# TBK1 restricts IRGQ-mediated autophagy

**DOI:** 10.64898/2026.01.07.698103

**Authors:** Uxia Gestal-Mato, Pauline Lascaux, Sergio Alejandro Poveda-Cuevas, Alberto Cristiani, Belinda Camp, Mahyar Aghapour, Aparna Viswanathan Ammanath, Ramachandra M Bhaskara, Ivan Dikic, Lina Herhaus

## Abstract

The autophagy-lysosome system directs the degradation of a wide variety of cytoplasmic cargo such as damaged organelles, protein aggregates, and invading pathogens. The autophagy receptor IRGQ harbors two distinct LIR domains, with LIR1 exhibiting high selectivity for GABARAPL2. Proteomic, biochemical, and high-throughput microscopy studies revealed that the IRGQ-GABARAPL2 complex functions as a hub for the interaction between hATG8s and the autophagy initiation machinery, promoting their lipidation and overall autophagic flux. The interaction of IRGQ with GABARAPL2 is regulated via TBK1. Upon TBK1 activation, GABARAPL2 is phosphorylated on S10, which disrupts IRGQ-GABARAPL2 complexation and therefore its interaction with the autophagy initiation machinery, resulting in a reduction of the autophagic flux of GABARAPL2 and IRGQ-cargo, without affecting bulk autophagy. These findings broaden IRGQ’s role in autophagy, identifying it as an interaction hub for autophagy initiation that is negatively regulated by TBK1.

## Introduction

Autophagy is a highly conserved, catabolic process that secures cellular homeostasis by recycling critical metabolites during starvation. Dysregulation of autophagy has been widely implicated in various pathophysiological processes such as aging, cancer, metabolic and neurodegenerative disorders, as well as cardiovascular and pulmonary diseases^1,2^. During the process of autophagy, a de-novo double-membraned vesicle termed the phagophore forms around demarcated cargo or bulk cytosol that will be degraded. The phagophore membrane arises from ATG9-positive vesicles, which serve as the initial membrane seed. ATG2-mediated lipid transport from the endoplasmic reticulum (ER) is required for expansion and subsequent phagophore formation^3,4^ and is governed by a multiprotein complex composed of ULK1/2, ATG13, FIP200, and ATG101^5–8^. Additionally, there are two systems involving ubiquitin-like proteins that contribute to the expansion of the phagophore and its subsequent fusion with the lysosome: the ATG12 conjugation pathway and the processing of the human ATG8 family (hATG8) by ATG7 and ATG3^9^.

The hATG8s are ubiquitin-like proteins that can be divided into two subfamilies: the LC3s (LC3A, LC3B, LC3C) and the GABARAPs (GABARAP, GABARAPL1, GABARAPL2). All members of this family are conjugated to phosphatidylethanolamine (PE) in the membrane of the growing phagophore (referred to as lipidation) by ATG7 (an E1-like enzyme), ATG3 (an E2-like enzyme), and the ATG12-ATG5-ATG16L1 complex (an E3-like enzyme)^10^. A prerequisite for PE conjugation is processing of pro-LC3 by the ATG4 protease. Closure of the phagophore results in the completed autophagosome that can fuse with lysosomes containing hydrolytic enzymes.

Based upon cargo being selectively delivered for degradation, autophagy has been classified into numerous subtypes, such as mitophagy, ER-phagy, or xenophagy^11^. Cargo selection in these pathways is often achieved via the binding of an autophagy receptor that harbors a consensus LC3-interacting region (LIR), a stretch of 4 amino acids to facilitate its direct binding to the LDS (LIR-docking site) of LC3 orthologs.

Autophagy receptors play a central role in linking specific cargo to the core autophagy machinery^12^. Thus, receptors serve as molecular adaptors that ensure selective cargo recognition and efficient sequestration within autophagosomes. In addition, some of them act as autophagy initiation hubs and can nucleate autophagosome formation by directly recruiting components of the core autophagy machinery, such as the ULK1 complex or ATG proteins, to the cargo site^13–15^. Receptors like NDP52 and OPTN have been demonstrated to interact with both cargo and core machinery, effectively coordinating cargo selection with autophagosome biogenesis. This dual role highlights how receptors actively shape the specificity and efficiency of selective autophagy responses, revealing them as dynamic organizers of selective autophagy, capable of modulating cargo degradation based on cellular context and signaling inputs.

We recently identified IRGQ as a novel receptor impacting the quality control of MHC class I molecules^16^. IRGQ interacts with GABARAPL2 and LC3B via two distinct conserved LIRs and is trafficked to lysosomes in an autophagy-dependent manner. The Tank1 binding kinase (TBK1) is a critical regulator of various forms of autophagy via the phosphorylation of autophagy receptors, promoting autophagic flux^17^. Additionally, our previous work demonstrated that TBK1 can also regulate LC3 orthologs to control autophagic flux^18^. Here, we present IRGQ as a hub for autophagy initiation that is restricted by TBK1. Through its unique dual LIR-binding, IRGQ serves as an interaction hub for the hATG8s and the autophagy initiation machinery, promoting their lipidation and flux. TBK1 activation results in phosphorylation of GABARAPL2 S10, causing the disruption of the hub and impairing the autophagic flux of LC3B, GABARAPL2, and IRGQ’s cargo molecule, HLA. All together, we uncover TBK1 as a negative regulator of IRGQ’s autophagic axis, showcasing the refinement of TBK1-regulated autophagic responses.

## Results

### IRGQ facilitates the interaction of hATG8s and the autophagy initiation machinery

Autophagy receptors harbor LIR domains crucial for interaction with hATG8s. IRGQ has two LIRs in its sequence, AlphaFold3 prediction shows LIR1 (aa186-189) facing the G-protein fold of its structure and LIR2 (aa421-424) in proximity to a disordered loop region (Fig. 1A). AlphaFold2-multimer modeling performed in a previous study^16^, suggested that out of the 6 hATG8s, GABARAPL2 was strongly preferred for the LIR1-LDS binding mode with 72% of the top models resulting in such complex (Fig. 1B, S1C).

**Figure 1:**
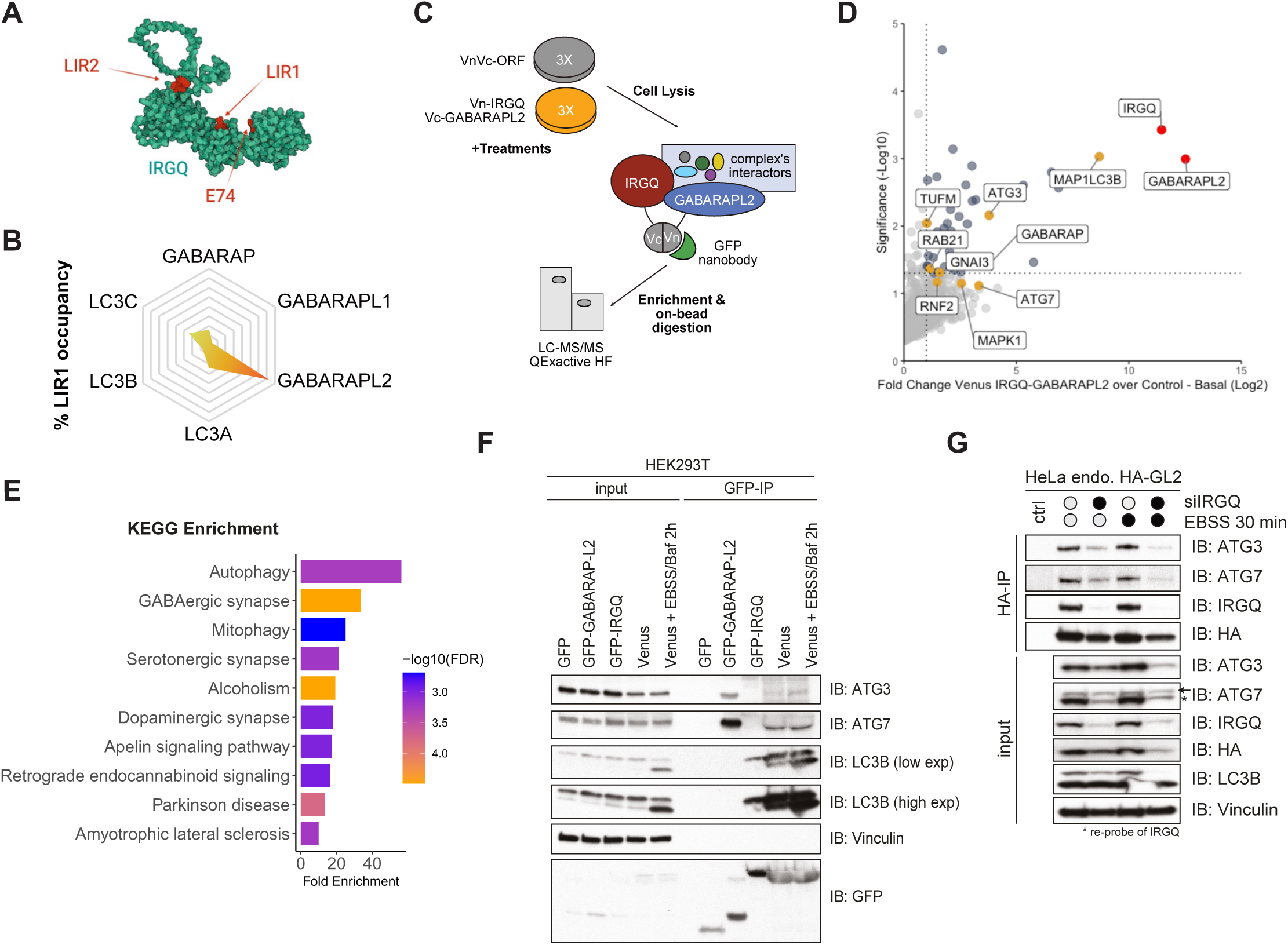
IRGQ facilitates the interaction of hATG8s and the autophagy initiation machinery. **(A)** IRGQ AlphaFold3 structure prediction highlighting interaction interfaces to hATG8s in red (E74R, LIR1, and LIR2). **(B)** Percentage of LIR1-LDS binding mode as the predicted complex between IRGQ-hATG8 in the 25 top-ranked AlphaFold2-multimer models from previous study^16^ **(C)** Experimental set-up for label free interactome studies of the IRGQ-GABARAPL2 complex. HEK293T cells were transfected with VnVc-ORF or Vn-IRGQ-Vc-GABARAPL2, lysates used for GFP-IPs and processed for mass spectrometry. Data was analyzed with MaxQuant and Perseus; n=3. **(D)** Volcano-plot representing the Student’s T-test difference from Vn-IRGQ-Vc-GABARAPL2 over VnVc-ORF IPs and -Log Student’s T-test p-value from Vn-IRGQ-Vc-GABARAPL2 over VnVc-ORF IPs in untreated conditions. The baits IRGQ and GABARAPL2 are marked in red, significant interaction partners in blue and autophagy-related proteins marked in yellow. **(E)** KEGG Enrichment Analysis of all significant interactors from Vn-IRGQ-Vc-GABARAPL2 over VnVc-ORF IPs in Basal conditions performed with ShinyGO 0.77. **(F)** SDS-PAGE and Western blot of GFP-IPs from HEK293T transfected with GFP-empty, GFP-GABARAPL2, GFP-IRGQ or Venus (Vn-IRGQ-Vc-GABARAPL2). **(G)** SDS-PAGE and Western blot of HA-IPs from endogenously tagged HA-GABARAPL2 cells upon treatment with IRGQ siRNA and/or EBSS starvation for 30 minutes.

Although there was high occupancy by LC3B for the LIR2-LDS models, there was no clear preference towards a specific hATG8 (Fig. S1A). The presence of two distinct LIR domains in the autophagy receptor IRGQ with one LIR motif having such specificity towards a single hATG8 is not known for any other autophagy receptor. To further study this phenomenon, we compared the N-terminal sequences of all hATG8s since the α-2 of GABARAPL2 is key for the interaction interface with IRGQ^16^. Sequence comparison shows that GABARAPL2 is the only hATG8 able to fit in the LIR1-G1 loop pocket of IRGQ, due to the presence of bulky residues in the other members of the family that would impede binding (Fig. S1B). GST pulldowns confirmed that the N-terminal region of GABARAPL2 is crucial for interaction with IRGQ (Fig. S1D).

To further dissect this striking evolutionary specificity, we queried the interactome of the IRGQ-GABARAPL2 complex with proteomics by using the Venus complementation affinity purification (BiCAP) method (Fig. 1C), which forms both in basal and autophagy-induced conditions (Fig. S1E). Immunoprecipitations using GFP-trap nanobodies in HEK293T cells revealed that LC3B, GABARAP, ATG3 and ATG7 are enriched in Venus-IPs versus control vectors, and that the binding to GABARAP and LC3B is further increased upon autophagy induction (Fig. 1D, S1F). KEGG Pathway Enrichment shows that interactors of the complex belong to pathways related to autophagy and membrane protein trafficking (Fig. 1E). These results were confirmed with co-immunoprecipitation using the Venus expressing cell lines, where GFP-GABARAPL2 binds endogenous ATG3 and ATG7, while GFP-IRGQ binds endogenous LC3B (Fig. 1F). Interestingly, when IRGQ and GABARAPL2 form a complex (Venus), they bind all three proteins (Fig. 1F). Conversely, the depletion of IRGQ from cells decreases the binding of endogenous GABARAPL2 to endogenous ATG3 and ATG7 (Fig. 1G). Due to both LIR motifs, IRGQ affects the co-precipitation of GABARAPL2, LC3B, and the autophagy initiation machinery. Taken together, the IRGQ-GABARAPL2 complex enables the recruitment of autophagy machinery and LC3B, acting as an autophagic hub.

### The IRGQ-GABARAPL2 autophagy hub promotes hATG8 lipidation

ATG3 (an E2-like enzyme) and ATG7 (an E1-like enzyme) mediate the conjugation of ATG8 proteins to phosphatidylethanolamine (PE), a key step in the autophagy pathway^10^. Since IRGQ affects the interaction between the hATG8s and the autophagy initiation machinery, we tested whether IRGQ assists in GABARAPL2 or LC3B lipidation. The overexpression of WT IRGQ results in increased LC3B (Fig. 2A, S2A) and GABARAPL2 lipidation (Fig. 2B). IRGQ mutants targeting the N-terminal region that mediates interaction with GABARAPL2 fail to promote GABARAPL2 lipidation despite comparable expression levels, demonstrating that lipidation depends on the integrity of the IRGQ-GABARAPL2 interaction rather than on IRGQ overexpression alone. Similarly, cells expressing the IRGQ-GABARAPL2 complex (Venus) exhibit increased LC3B lipidation at early time points of EBSS induction, compared to empty vector (Fig. 2C, S2B). Interestingly, overexpressing GABARAPL2 alone seemingly increases the unlipidated LC3B, possibly due to a sequestration of ATG3 and ATG7 by GABARAPL2 (Fig. 2C).

**Figure 2:**
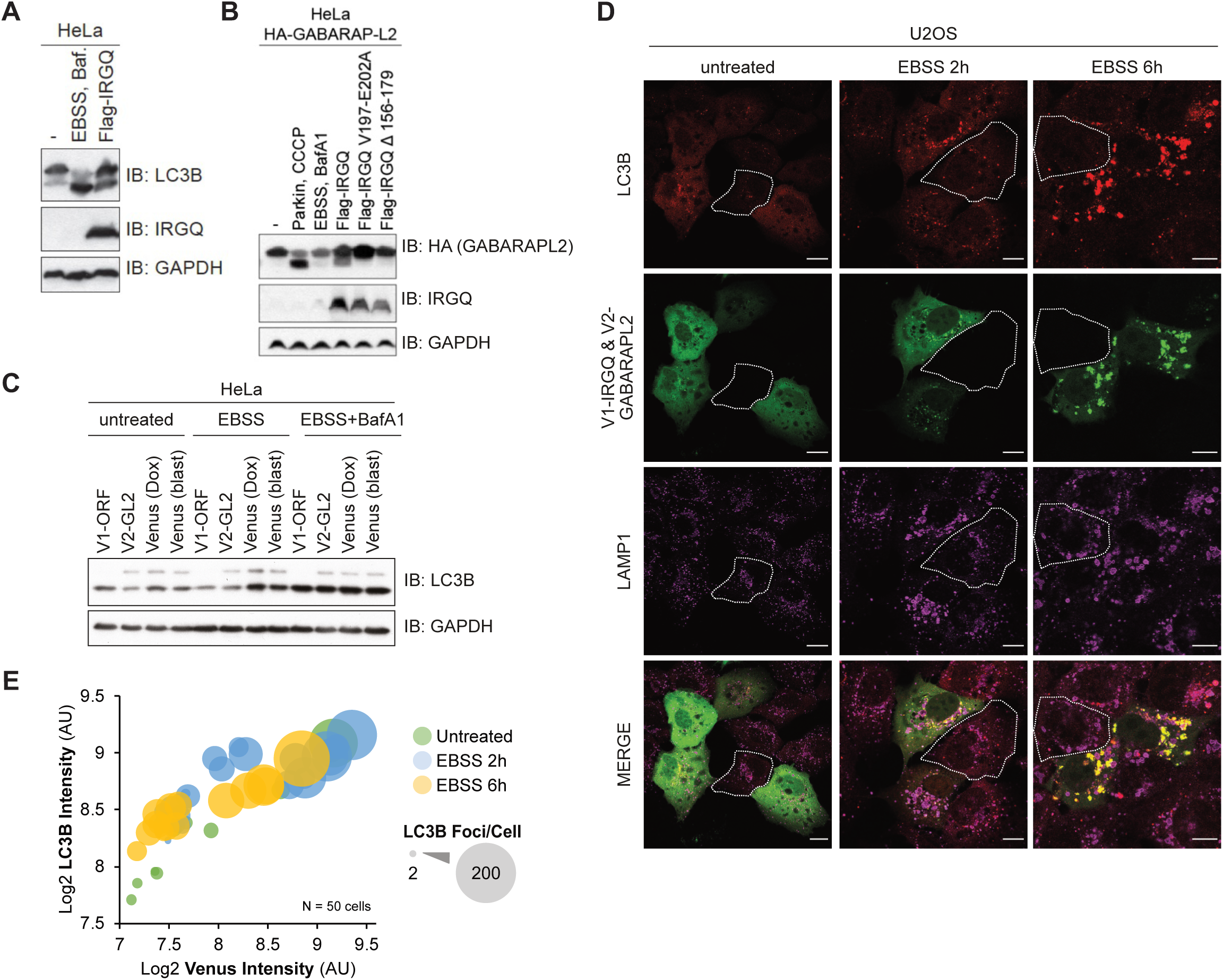
The IRGQ-GABARAPL2 complex promotes hATG8 lipidation. **(A)** SDS-PAGE and Western blot of HeLa cell lysates expressing transfected Flag-IRGQ or treated with EBSS, 200 nM Bafilomycin A1 (3 h). **(B)** SDS-PAGE and Western blot of endogenously tagged HA-GABARAPL2 HeLa cell lysates with transfected Flag-IRGQ WT or mutants, treated with EBSS, 200 nM Bafilomycin A1 (3 h) or treated for CCCP 40 µM (3h). **(C)** SDS-PAGE and Western blot of HeLa cell lysates stably expressing V1-ORF, V2-GABARAPL2, V1-IRGQ or V1-IRGQ & V2-GABARAPL2 (Venus). Cells were either left untreated or treated with EBSS or EBSS and BafA1 (200 nM) for 3 hours. **(D)** Immunofluorescence of HeLa cells, stably expressing V1-IRGQ and V2-GABARAP-L2. Autophagy was induced by the addition of EBSS for 2 hours or 6 hours. In addition, cells were left untreated. Fixed cells were probed with an endogenous LC3B or LAMP1 antibody. Scale bar: 20 µm. **(E)** Scatter plot of ImageJ quantification from (D). Single cells were annotated as ROIs and intensities and puncta for LC3B and Venus were measured. The boxed cells are non-transfected control cells for comparison to the transfected cells. RawIntDen values are plotted as Log2. Colors indicate the treatment and size of bubbles indicate the number of LC3B puncta per cell; n=50 cells.

Upon autophagy induction, hATG8s are lipidated and decorate autophagosomal membranes. In particular, LC3-II foci formation is often used as a measurement for autophagic flux^19,20^. Thus, we used immunofluorescence studies to monitor the effect of the IRGQ-GABARAPL2 complex on LC3B lipidation. Confocal images of cells expressing the IRGQ-GABARAPL2 Venus complex suggested that expressing-cells had a higher level of LC3B staining intensity and puncta formation in comparison to non-expressing cells, highlighted in white (Fig. 2D). Image quantification confirmed this observation, as Venus intensity positively correlated with LC3B intensity and more LC3B foci were present in those cells where the levels of the IRGQ-GABARAPL2 complex was higher (Fig. 2E). Analysis of LAMP1 staining reveals similar results regarding the number of foci, although intensity does not vary as much in relation to Venus expression, compared to LC3B (Fig. S2C). To directly assess autophagy flux and exclude the possibility that increased LC3B lipidation and foci reflect a block in autophagy, we analyzed p62 turnover by immunoblotting (Fig. S2D). As expected, p62 levels were reduced upon EBSS treatment, confirming active autophagy flux. Importantly, upon Venus-based co-overexpression of IRGQ and GABARAPL2, p62 degradation was further enhanced, indicating increased autophagic flux rather than impaired turnover.

### IRGQ-GABARAPL2 complex stability is disrupted upon GABARAPL2 S10D mutation

The interaction interface between the two proteins shows E74 of IRGQ, a key residue for the complex interaction^16^, bridging to S10 of GABARAPL2 (Fig. 3A). Notably, this residue is heavily conserved among species (Fig. S3A). Our previous research showed that TBK1 phosphorylates members of the ATG8 family on several residues, GABARAPL2-S10 among them^18^. Mutational analysis in a TBK1 *in vitro* kinase assay reveals that S10 is the major TBK1-mediated phosphorylation site, since the S10A mutant abrogates almost completely the phosphorylation of the protein (Fig. 3B). Correspondingly, *in cellulo*, S10 is the most prominent GABARAPL2 phosphorylation site. Cell extracts of HEK293T cells overexpressing TBK1 and GABARAPL2 WT or non phosphorylatable alanine mutants were analyzed on a PhosTag gel, where phosphorylated proteins are retained and are represented by an upwards shift. Mutation of GABARAPL2 S10A results in a loss of phosphorylated protein (Fig. S3B). Similarly, mutation of TBK1 K38A (kinase dead mutant) abolished GABARAPL2 phosphorylation (Fig. S3C).

**Figure 3:**
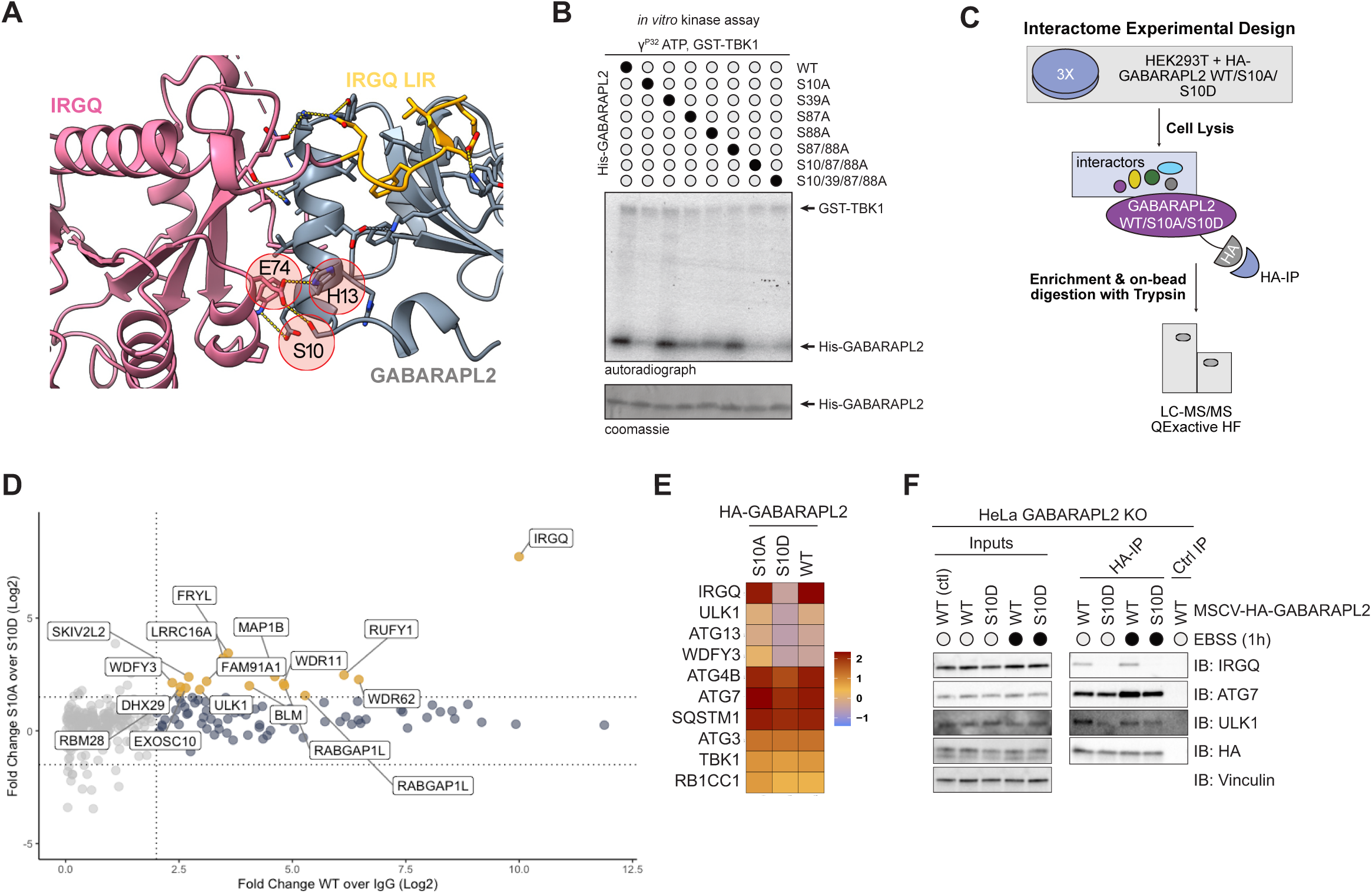
IRGQ-GABARAPL2 complex stability is disrupted upon GABARAPL2 S10D mutation. **(A)** Structure of IRGQ N-terminal domain (pink) and IRGQ LIR1 185-190 (yellow) in complex with GABARAPL2 (gray). GABARAPL2 α-helix 2 makes contacts within a pocket formed by IRGQ’s switch I (partially disordered), switch II and the linker between LIR1 and N-terminal G-domain. Highlighted are key residues for this additional binding interface (E74 from IRGQ, H13 and S10 from GABARAPL2). **(B)** Coomassie stain and autoradiography of SDS-PAGE after an *in vitro* kinase assay with GST-TBK1, His-GABARAP-L2 WT and mutant proteins as substrates. **(C)** Experimental set-up for label free interactome studies of the GABARAPL2 S10 mutants. HEK293T cells were transfected with HA-GABARAPL2 WT/S10A or S10D mutants, lysates used for HA-IPs and processed for mass spectrometry. Data was analyzed with MaxQuant and Perseus; n=3. **(D)** Scatter plot representing the Student’s T-test difference from HA-GABARAPL2 over empty control IPs and the Student’s T-test difference from HA-GABARAPL2 S10A over HA-GABARAPL2 S10D IPs. Significant and common interaction partners are marked in blue and interaction partners showing preference of binding towards S10A mutant are marked in yellow (Fold Change S10A over S10D ≥ 2). **(E)** Heatmap of selected autophagy-related proteins. Color scale represents the z-score of the average LFQ Intensity. **(F)** SDS-PAGE and Western blot of HA-IPs from GABARAPL2 KO cells reconstituted with HA-GABARAPL2 WT of S10D mutant (with or without EBSS treatment for 1 hour).

To understand the relevance of this phosphorylation we transfected HEK293T cells with different constructs of GABARAPL2, performed pulldowns of the mutants and used proteomics to analyze the variation in their interactomes (Fig. 3C). GABARAPL2 WT interacts with known partners belonging to biological processes such as macroautophagy, selective autophagy and intracellular transport^21,22^, as shown with Gene Ontology Enrichment analysis (Fig. S3D). Comparison of the three pulldowns highlights that the phospho-mimicking mutant S10D loses affinity to several proteins compared to WT and S10A, which share the majority of interactors (Fig. 3D). The interaction of GABARAPL2 S10D to IRGQ, ULK1, ATG13, ATG4B and WDFY3 is significantly decreased, together with a small reduction in binding to ATG7, and p62/SQSTM1 (Fig. 3E). The loss in affinity to IRGQ was expected due to the previously described interaction mode of GABARAPL2 N-terminal to IRGQ, and was confirmed by WB of different pulldowns (Fig. S3E, S3F). To validate the MS findings on the other autophagy-related proteins, we made use of reconstituted cell lines from GABARAPL2 KO cells where the HA-GABARAPL2 WT or mutant genes were reintroduced with lentiviral transductions (Fig. S3G). HA-GABARAPL2 WT interacts with IRGQ, ATG7, and ULK1 under both basal and starvation conditions, whereas the HA-GABARAPL2 S10D mutant shows a pronounced reduction in IRGQ binding, with modest effects on ATG7 and ULK1 interactions (Fig. 3F). This selective loss of IRGQ engagement is consistent with our interactome analysis (Fig. 3E) and with the reduced association of these factors observed upon IRGQ knockdown (Fig. 1G). Autophagy-related proteins such as ULK1 typically bind ATG8 family members through LIR motifs that interact with the LDS ^23,24^. Ser10 in GABARAPL2 is located at a considerable distance from the LDS (Fig. S3H) and its phosphorylation selectively impairs IRGQ binding. Because IRGQ likely stabilizes or promotes the assembly of complexes with ATG7 and ULK1, the weakened IRGQ-GABARAPL2 interaction in the S10D mutant may indirectly account for the subtle changes observed with these autophagy-initiation factors. This supports a model in which IRGQ functions as an interaction hub coordinating GABARAPL2 with components of the autophagy-initiation machinery.

To provide structural insights into how TBK1-dependent phosphorylation modulates this selective interaction, we performed molecular modelling of phosphorylated GABARAPL2 in the context of the IRGQ-ATG8 interface and additional ATG8-dependent assemblies. These models position Ser10 within the IRGQ-binding surface of GABARAPL2, such that the phosphate group (or the S10D phosphomimetic variant) perturbs the local electrostatic and steric environment at the IRGQ interface, thereby weakening IRGQ engagement. Importantly, our modeling showed that Ser10 of GABARAPL2 is spatially separated from the canonical LDS (LIR docking site), consistent with the experimental observation that LDS-mediated interactions of GABARAPL2 with other ATG8 interactors are largely preserved. Together, these data (Fig. S4A-H) point to a mechanism where TBK1 phosphorylation selectively destabilizes the IRGQ-GABARAPL2 complex while preserving broader ATG8 interactomes.

Consistent with this model, AlphaFold2-multimer predictions indicate that IRGQ can organize into higher-order assemblies with ATG8 proteins and the lipidation machinery. In predicted IRGQ-GABARAPL2-ATG7 complex models, ATG7 consistently engage with the canonical LDS of the GABARAPL2 interface (> 70%). By contrast, alternative configurations in which IRGQ and ATG7 compete for or exchange interfaces on GABARAPL2 are comparatively rare (< 20%) (Fig. S5A). Extending this modeling to a four-component system, IRGQ-GABARAPL2-LC3B-ATG7 predominantly (> 70%) place ATG7 on GABARAPL2 LDS while IRGQ engages with LC3B, with only a minor fraction showing swapped binding mode (ATG7 bound to LC3B and IRGQ bound to GABARAPL2) (Fig. S5B). Together, these models support our experimental data and indicate a scaffold mechanism where IRGQ, through its dual-LIR architecture, can simultaneously engage GABARAPL2 and LC3B, thereby promoting proximity to ATG7 and other ATG8-dependent factors without requiring direct IRGQ-ATG7 binding. Rather than directly abrogating LDS-mediated interactions, phosphorylation of GABARAPL2 at S10 by TBK1 selectively disrupts the IRGQ-GABARAPL2 interface of these scaffold complexes, indirectly destabilizing the assembly to reduce the overall recruitment efficiency of ATG8-associated machinery.

### TBK1 phosphorylation of GABARAPL2-S10 restricts the IRGQ-GABARAPL2 autophagy hub

TBK1 has been demonstrated to play a critical role in selective autophagy via the regulation of a variety of autophagy proteins^25,26^. In order to test if TBK1 can phosphorylate GABARAPL2 on S10 in cells, we raised an antibody that specifically detects GABARAPL2 pS10 (Fig. S6A). TBK1 kinase activity can be stimulated under conditions that elicit selective autophagy (such as mitophagy or xenophagy). Induction of mitophagy as well as xenophagy and IFN treatment strongly induced endogenous GABARAPL2 S10 phosphorylation, especially of the lipidated form (Fig. 4A, S6B). However, the induction of starvation-induced bulk autophagy, which does not activate TBK1, did not lead to a phosphorylation of GABARAPL2 (Fig. 4A, S6B).

**Figure 4:**
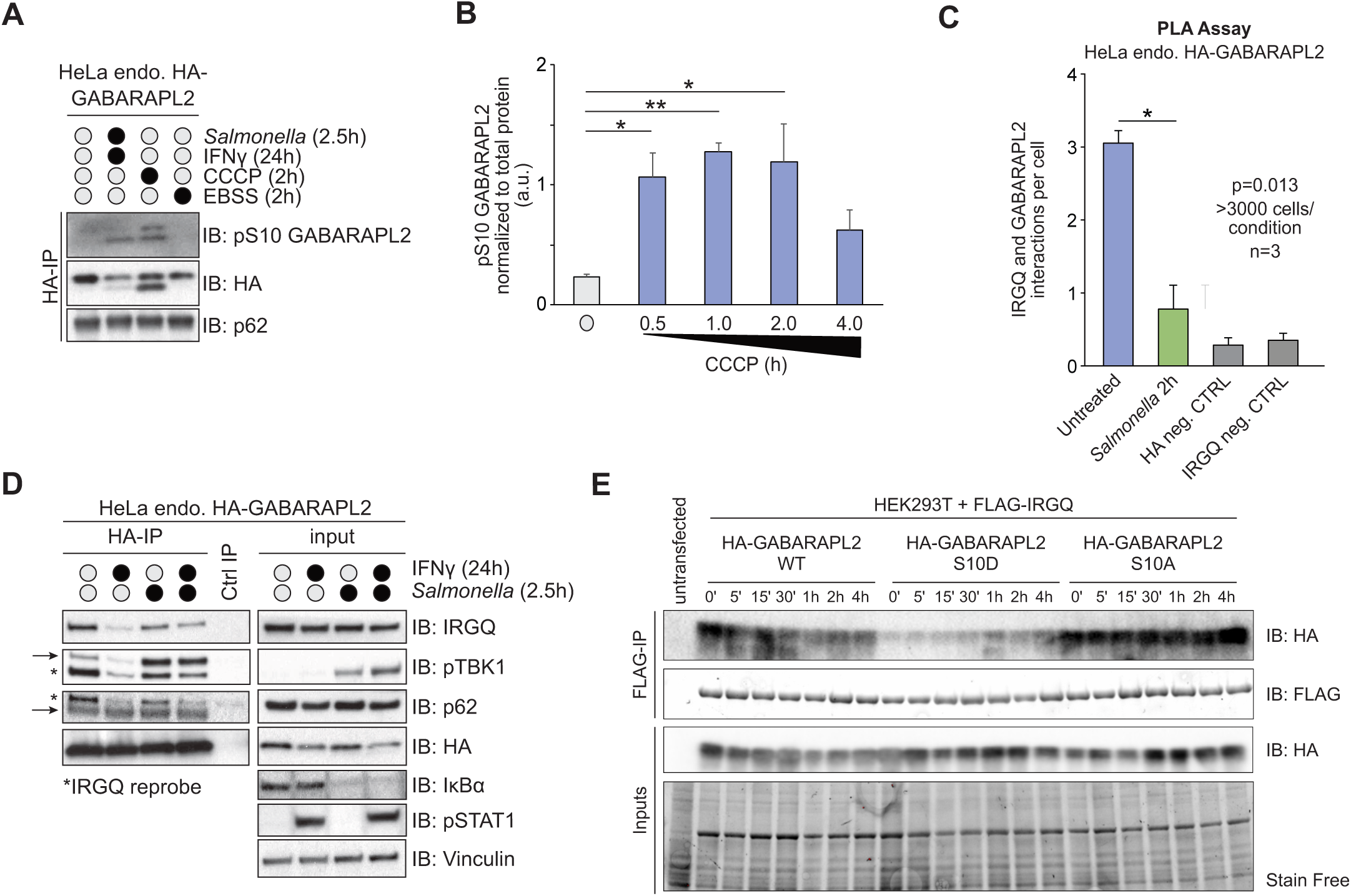
TBK1 phosphorylation of GABARAPL2-S10 restricts the IRGQ-GABARAPL2 autophagy complex. **(A)** SDS-PAGE and Western blot of HA-IPs from endogenously tagged HA-GABARAPL2 WT HeLa cells after treatment with CCCP (2h), EBSS (2h) and IFNy (24h) plus *Salmonella* infection (2.5 h). **(B)** ImageJ quantification from (S4E) of pS10 GABARAPL2, normalized to total GABARAPL2 protein. Data are presented as the mean with error bars indicating the s.d. Statistical significance of differences between experimental groups was assessed with Student’s T-test. Differences with p < 0.05 are annotated as * and p < 0.01 are annotated as **; n = 3. **(C)** Yokogawa CQ1 quantification of average Duolink PLA signal from endogenous GABARAP-L2 and endogenous IRGQ. HeLa endogenous HA-GABARAPL2 cells were infected with *Salmonella* for 2 hours and fixed cells were probed with the Duolink in situ PLA assay. HA and IRGQ only antibodies were used as negative controls to determine the background. Data are presented as the mean with error bars indicating the s.d. Statistical significance of differences between experimental groups was assessed with Student’s T-test. Differences with p < 0.05 are annotated as *; n=3; >3000 cells/condition. **(D)** SDS-PAGE and Western blot of HeLa endogenous HA-GABARAP-L2 cells that were treated with 10 ng/ml IFNγ (24 h) and infected with WT *Salmonella* (2.5 h). Cell lysates were subjected to HA-IPs or IgG control IP. **(E)** SDS-PAGE and Western blot of HEK293T exogenously expressing Flag-IRGQ and HA-GABARAPL2 WT or S10 mutants. Cells were treated with 40uM CCCP at specified timepoints. Cell lysates were subjected to FLAG-IPs.

The appearance of this modification was dependent on the presence of TBK1 and specifically its kinase activity, as knockdown or chemical inhibition with MRT67307 or BX795 blocked phosphorylation of S10 GABARAPL2 (Fig. S6C, S6D). Intriguingly, TBK1 preferentially phosphorylated PE-conjugated His-GABARAPL2 as seen in an *in vitro* kinase assay (Fig. S6E). To more closely delineate the kinetics of this phosphorylation, we induced mitophagy and measured GABARAPL2 pS10 at various time points. GABARAPL2 pS10 peaks at 1h after treatment and coincides with maximal TBK1 activation (Fig. 4B, S6F). Similarly, infection with *Salmonella* to trigger TBK1 activation results in phosphorylation of S10 GABARAPL2, with similar kinetics as mitophagy induction (Fig. S6G). Notably, GABARAPL2 pS10 can also be detected in human samples from HBV-infection derived tumors (Fig. S6H).

Since phospho-mimicking mutations of GABARAPL2 S10 restricted its interaction with IRGQ, we confirmed that TBK1 activation in cells causing phosphorylation of S10 on GABARAPL2 had the same effect on its binding to endogenous IRGQ. Indeed, TBK1 stimulation upon xenophagy induction causes a significant reduction in IRGQ-GABARAPL2 endogenous foci measured with proximity ligation assays (PLA) (Fig. 4C, S6I, S6J). Furthermore, co-immunoprecipitation of endogenous GABARAPL2 with IRGQ is substantially restricted upon TBK1 stimulation as seen by Western Blot (Fig. 4D, S6K). To further validate the effect of TBK1 on the regulation of the IRGQ-GABARAPL2 hub, HEK293T cells were co-transfected with Flag-IRGQ and HA-GABARAPL2 WT, S10D or S10A mutant constructs and subjected to a time course treatment of CCCP to trigger TBK1 activation. In basal conditions, HA-GABARAPL2-WT interacts strongly with Flag-IRGQ but the complex is disrupted upon CCCP treatment, with the lowest interaction signal coinciding with the highest peak of TBK1-mediated S10 phosphorylation of GABARAPL2 at 1 hour after treatment (Fig. 4B, E). In contrast, HA-GABARAPL2-S10A interaction with Flag-IRGQ is unaffected upon TBK1 activation, showing a slight increase at the 4 hours timepoint. As expected, HA-GABARAPL2-S10D mutant does not interact with Flag-IRGQ even in basal conditions (Fig. 4E). Hence, GABARAPL2 S10 is specifically modified by TBK1, once it is activated and causes IRGQ-GABARAPL2 complex disruption.

### TBK1 is a negative regulator of IRGQ-mediated autophagy

The IRGQ-GABARAPL2 hub interacts with the autophagy initiation machinery and facilitates the lipidation of LC3B. Upon TBK1 activation, GABARAPL2 is phosphorylated on S10 causing the dissociation from IRGQ and autophagy initiation proteins (Fig. 5A). Therefore, we tested if this regulatory modification would also impact LC3B lipidation or flux. To assess whether phosphorylation of GABARAPL2 at Ser10 impacts autophagy progression, we performed a high-throughput microscopy time-course assay coupled to automated image analysis in HA-GABARAPL2-WT, S10A and S10D reconstituted cells. We quantified LC3-to-lysosome flux in cells expressing GABARAPL2 WT, S10A, or S10D (Fig. 5B). Each data point represents the normalized mean LC3-to-lysosome ratio per well and per biological replicate, normalized within each repeat to the untreated WT mean. Conditions included starvation (EBSS, 1-4 h), mitochondrial depolarization (40 µM CCCP, 1-4 h), and lysosomal inhibition (200 nM BAF for 4 h, ± EBSS or CCCP). Across these conditions, LC3-to-lysosome flux was not significantly altered by changes in GABARAPL2-S10 phosphorylation, indicating that S10D does not measurably impair inducible bulk autophagy flux. Together, these data support the conclusion that the primary impact of the S10D mutation is on IRGQ engagement and the basal organization of ATG8-positive structures, rather than on global autophagy induction or lysosomal delivery of LC3 under stress.

**Figure 5:**
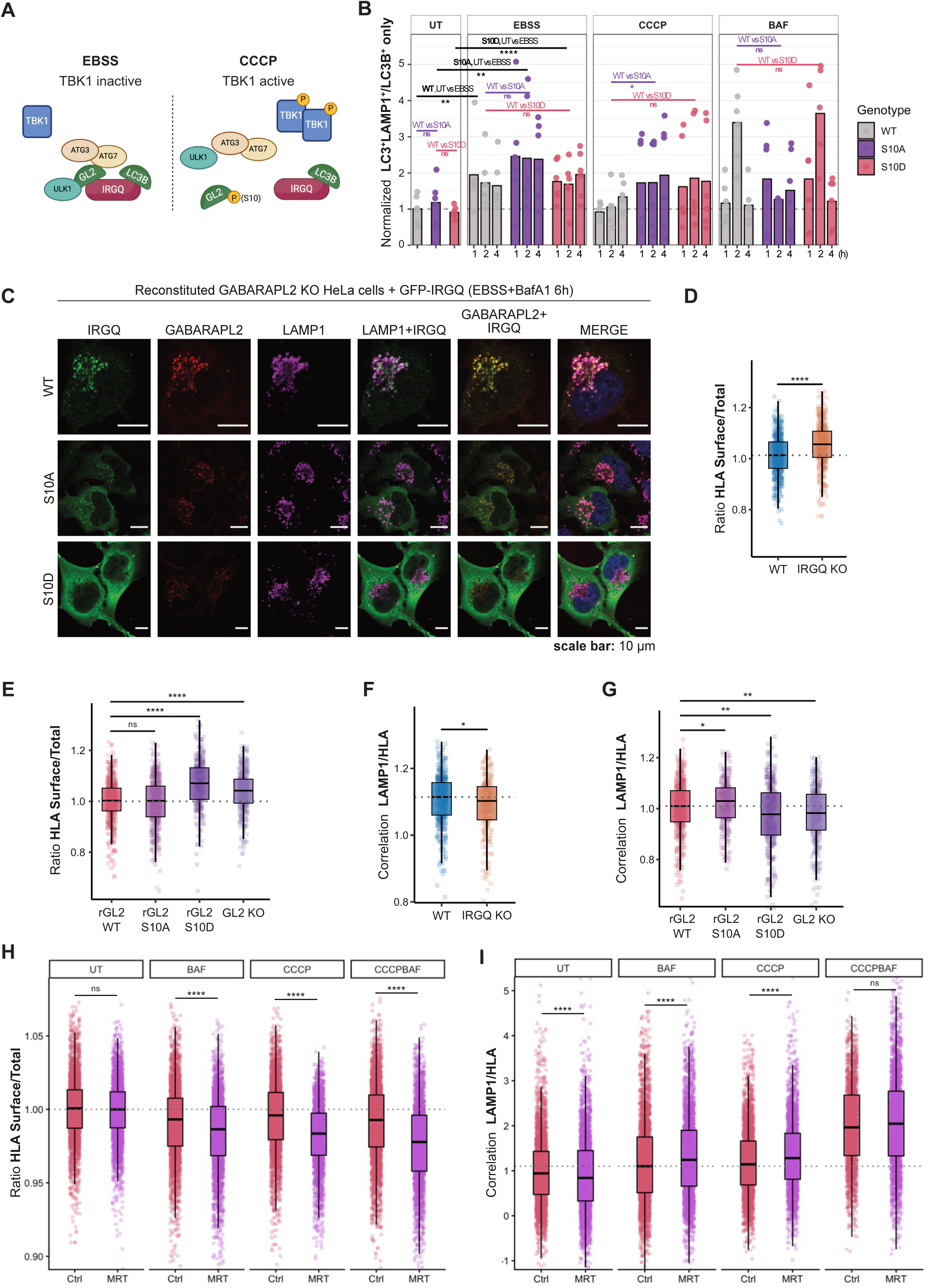
TBK1 is a negative regulator of IRGQ-mediated autophagy. **(A)** Model of IRGQ autophagy hub in basal conditions or upon TBK1 activation. Schematic was done with BioRender. **(B)** LC3-to-lysosome flux under stress is not changed when GABARAPL2 S10 phosphorylation is altered. Quantification of LC3-to-lysosome flux in GABARAPL2 WT, S10A, and S10D cells. Each point represents the normalized mean ratio per well and per biological repeat (three independent experiments, two wells per condition). Values were normalized to the untreated (UT) WT mean within each repeat. Conditions represent starvation (EBSS, 1-4 h), mitochondrial depolarization (40 µM CCCP, 1-4 h), or lysosomal inhibition (200 nM BAF for 4 h; ± EBSS or CCCP). Grouped one-sample t-tests compared combined EBSS (E1-E4), CCCP (C1-C4), and BAF (BAF, EBSS + BAF, CCCP + BAF) groups to UT (= 1). **** p < 0.0001; *** p < 0.001; ** p < 0.01; * p < 0.05; n.s., not significant. **(C)** Immunofluorescence of HeLa GABARAPL2 KO cells reconstituted with WT, S10A or S10D HA-GABARAPL2 and overexpressed GFP-IRGQ. Autophagy was induced by the addition of EBSS for 6 hours and Bafilomycin A1 (200 nM). Fixed cells were probed with endogenous LAMP1 and HA antibodies. Scale bar: 10 µm. **(D,E)** Boxplots of normalized LAMP1/HLA correlation. Automated quantification done with CellProfiler. All tests are two-sided Wilcoxon rank-sum comparisons; p values were adjusted by the Benjamini-Hochberg FDR procedure. Significance codes: ns ≥ 0.05, * < 0.05, ** < 0.01, *** < 0.001, **** < 0.0001. Four biological replicates were analyzed.; n=>100 cells/replicate. **(F,G)** Boxplots of normalized HLA Rim/Total cell intensity per condition after 4 hours of EBSS and Bafilomycin A1 (200 nM) treatment. Values were normalized to the mean of WT within each replicate. Each point represents a single; boxes show the interquartile range with median lines. Automated quantification done with CellProfiler. All tests are two-sided Wilcoxon rank-sum comparisons; p values were adjusted by the Benjamini-Hochberg FDR procedure. Significance codes: ns ≥ 0.05, * < 0.05, ** < 0.01, *** < 0.001, **** < 0.0001. Four biological replicates were analyzed.; n = >100 cells/replicate. **(H)** Quantification of HLA-to-lysosome flux and HLA presentation at the cell surface in HeLa WT cells. Each point represents the normalized mean ratio per well and per biological repeat (two independent experiments, three wells per condition). Values were normalized to the untreated (UT) mean within each repeat. Conditions represent mitochondrial depolarization (40 µM CCCP, 4 h), or lysosomal inhibition (200 nM BAF for 4 h) with or without TBK1 inhibition (MRT67307 5 μM). Wilcoxon rank-sum test on single-cell values: **** p < 0.0001; *** p < 0.001; ** p < 0.01; * p < 0.05; n.s., not significant.

In contrast to LC3B flux, GABARAPL2 autophagic flux itself was altered by the phospho-mimicking mutant. To monitor GABARAPL2 trafficking directly, we generated U2OS reporter cell lines expressing mCherry-GFP-tagged GABARAPL2 WT, S10A, or S10D and quantified flux using automated high-throughput imaging (Fig. S7A,B) and FACS (Fig. S7C). Upon EBSS starvation, WT and S10A cells showed an increase in GABARAPL2 flux, and this response was blocked by BafA1, consistent with lysosome-dependent turnover. By contrast, S10D cells exhibited lower basal GABARAPL2 flux, and starvation failed to further enhance flux (Fig. S7A). Across both readouts, S10A closely phenocopied WT, whereas S10D displayed a selective flux defect consistent with a constitutive phospho-mimetic state. These data support a model in which TBK1-mediated phosphorylation of GABARAPL2 at Ser10 functions as a conditional negative regulatory (“stop”) signal, terminating IRGQ-GABARAPL2 engagement and limiting lysosomal trafficking once the relevant step of the pathway has been completed.

To validate that the S10D mutation disrupts delivery of the GABARAPL2-IRGQ complex to lysosomes, we performed immunofluorescence analyses. Under starvation-induced autophagy, GFP-IRGQ formed punctate structures that colocalized with GABARAPL2 and LAMP1 in reconstituted GABARAPL2 WT cells (Fig. 5C). Similarly, GFP-IRGQ puncta colocalizing with LC3B and LAMP1 were observed in GABARAPL2 S10A cells (Fig. 5C). In contrast, in S10D cells GFP-IRGQ remained diffusely cytosolic and failed to form puncta (Fig. 5C), underscoring that a tightly regulated Ser10 phosphorylation state is required for IRGQ-dependent autophagy. In line with this model, WT GABARAPL2 is expected to be largely unphosphorylated under basal conditions and phosphorylation to be transient and context-dependent, providing a mechanistic explanation for why the non-phosphorylatable S10A mutant often resembles WT.

Recently, we described IRGQ as an autophagy receptor that functions in the quality control of ‘non-conformational’ HLA molecules^16^. Naturally, we tested if this function of IRGQ was also negatively regulated by the TBK1-mediated phosphorylation of GABARAPL2, since flux of IRGQ to the lysosome is impaired upon GABARAPL2 depletion^16^. Immunofluorescence analysis and quantification with CellProfiler confirmed that HLA (IRGQ’s cargo) was accumulated at the cell surface of S10D mutants, mimicking the phenotype of IRGQ KO cells (Fig. 5D,E, S7D). This was seemingly a cause of impaired lysosomal delivery of HLA molecules observed in these cells (Fig. 5F,G, S7D). In addition, we confirmed that pharmacological inhibition of TBK1 using MRT67307 further enhances lysosomal delivery of HLA molecules (Fig. 5H). Specifically, we quantified HLA-to-lysosome flux under stress conditions in the presence of MRT67307 and observed increased HLA accumulation in lysosomal compartments, consistent with our findings in Fig. 5D-G. Together, these results corroborate that TBK1 activity restrains IRGQ-dependent trafficking of HLA to lysosomes, and that TBK1 inhibition promotes HLA turnover via the lysosomal pathway under stress conditions.

To determine whether GABARAPL2-S10 phosphorylation broadly affects autophagy or selectively impacts IRGQ-dependent pathways, we assessed p62 turnover as a readout of bulk and parallel selective autophagy fluxes. We quantified p62-to-lysosome flux in cells expressing GABARAPL2 WT, S10A, or S10D under starvation (EBSS), mitochondrial depolarization (CCCP), and lysosomal inhibition (BAF) conditions (Fig. S7D). p62 degradation was comparable between WT, S10A, and S10D cells, indicating that p62-dependent autophagy flux is independent of GABARAPL2-S10 phosphorylation. These data demonstrate that S10 phosphorylation does not impair bulk autophagy (consistent with no impact on LC3B flux Fig. 5B), but instead selectively affects IRGQ-dependent cargo turnover, exemplified by MHC-I degradation (Fig. 5H).

In short, disruption of the IRGQ-GABARAPL2 autophagic hub by TBK1 stimulation as well as phospho-mimicking mutations of GABARAPL2 S10 affected the flux of GABARAPL2 and IRGQ’s cargo molecule HLA, but not bulk autophagy, revealing TBK1 as a negative regulator of IRGQ’s autophagic axis.

## Discussion

Autophagy is a tightly regulated catabolic pathway critical for cellular homeostasis, with selective autophagy receptors serving as key coordinators between cargo recognition and the core autophagy machinery^27^. Our study identifies the IRGQ-GABARAPL2 complex as a central organizer of autophagy initiation, extending the functional repertoire of IRGQ beyond cargo recognition to active orchestration of GABARAPL2 lipidation and flux. While IRGQ was previously characterized as a receptor for misfolded MHC class I molecules^16^, we now demonstrate that it additionally functions as a scaffold to promote autophagy initiation and is subject to regulation by TBK1.

We show that IRGQ binds to both GABARAPL2 and LC3B via two conserved LIR motifs, supporting its role as a dual hATG8 interactor. This dual engagement enables IRGQ to promote the recruitment and lipidation of GABARAPL2 and LC3B, thereby driving autophagosome formation. The importance of this interaction is underscored by the finding that IRGQ localizes to lysosomes in an autophagy-dependent manner, suggesting efficient flux under basal conditions.

Unexpectedly, we uncover that TBK1, which is widely recognized as a positive regulator of autophagy^28,29^, can act as a negative regulator in this context. TBK1 activation leads to phosphorylation of GABARAPL2 at Ser10, disrupting its interaction with IRGQ and thereby disassembling the complex. This in turn impairs the flux of GABARAPL2 and IRGQ’s cargo, misfolded MHC-I, to lysosomes, without affecting bulk autophagy. These findings reveal a previously unrecognized mechanism by which TBK1 fine-tunes autophagic responses through spatial and temporal control of hub assembly.

Importantly, this regulatory axis challenges the prevailing view of TBK1 as a uniformly pro-autophagic kinase. While TBK1-mediated phosphorylation of autophagy receptors is often associated with enhanced flux^30^, our data show that it can also suppress selective autophagy by dismantling key protein-protein interactions. Thus, the functional outcome of TBK1 activity appears to be highly context-dependent, varying with the identity of the autophagy receptor and cargo.

In this context, we propose that TBK1-mediated phosphorylation of GABARAPL2 at Ser10 acts as a negative regulatory termination signal. Specifically, phosphorylation at S10 disrupts IRGQ–GABARAPL2 engagement and thereby limits further trafficking of GABARAPL2-containing complexes to lysosomes once the relevant step of the pathway has been executed. Such a mechanism implies that S10 phosphorylation is transient and conditional, which provides a straightforward explanation for why the non-phosphorylatable S10A mutant often phenocopies WT: under basal conditions GABARAPL2 is expected to be largely unphosphorylated, and thus WT and S10A behave similarly. By contrast, S10D mimics constitutive phosphorylation, enforcing a persistent “off” state that blocks IRGQ binding and downstream lysosomal targeting, thereby revealing the importance of regulated disengagement rather than continuous interaction. AMPK is a well-established energy sensor and a key regulator of autophagy, traditionally recognized for its role in promoting autophagy under conditions of energy stress. However, recent findings underscore a more nuanced function of AMPK, revealing its dual role as both a promoter and inhibitor of autophagy depending on the cellular context and regulatory interactions^31^. This duality parallels our findings, where we now demonstrate that TBK1, similarly known for its autophagy-promoting activities, can also function as a negative regulator by inhibiting the binding of GABARAPL2 to IRGQ. These insights highlight the context-dependent and multifaceted roles of kinases in autophagy regulation.

Our work also provides mechanistic insights into the control of MHC class I quality via autophagy. By demonstrating that IRGQ facilitates not only cargo recognition but also initiation of autophagy, we suggest that certain receptors act as integrators of cargo selection and machinery assembly. This positions IRGQ at the nexus of immune regulation and proteostasis, with implications for understanding immune evasion, antigen presentation, and inflammatory signaling.

In summary, we propose a model in which the IRGQ-GABARAPL2 complex functions as a scaffolding complex that nucleates the autophagy initiation complex through its dual hATG8 interactions and the recruitment of the autophagy-initiation machinery, and whose activity is negatively regulated by TBK1-mediated phosphorylation of GABARAPL2. These findings expand our understanding of selective autophagy regulation and reveal new layers of control by kinase signaling networks.

## Methods

### Expression constructs

Expression constructs of indicated proteins were cloned into indicated vectors using PCR or the gateway system. Site-directed mutagenesis was performed by PCR to introduce desired amino acid substitutions. All expression constructs were sequenced by Seqlab.

### Cell culture

HEK293T, U2OS and HeLa cells were cultured in Dulbecco’s modified Eagle’s medium (DMEM; Gibco) supplemented with 10% fetal bovine serum (FBS), 1% penicillin/streptomycin and maintained at 37°C in a humidified atmosphere with 5% CO_2_. For Salmonella infections penicillin/streptomycin was omitted from the media. HeLa WT endogenous HA-GABARAPL2 cells were generated using CRISPR/Cas9 technology and kindly provided by the Behrends laboratory (LMU München, Germany). Lentiviral stable cell lines were generated as previously described ^32^. The guide RNAs used for the CRISPR/Cas 9 KO cells are: IRGQ #1: TTTGTGCTACCGGCGAACTG, IRGQ #2: GAATGCACTCAGTAAGGGAA, IRGQ #3: CGTGAGGCCTTTGAGACCGG, GABARAPL2 #1: GTCGAGCGAAATATCCCGACA, GABARAPL2 #2: GTCCCACAGAACACAGATGCG, GABARAPL2 #3: GGTTCCATCTGATATCACTG, and were cloned into Lentiviral vectors containing CAS9: pLenti-Puro or pLenti-Neo (as described previously ^32^). After viral infection cells were selected in media containing 2 µg/ml of Puromycin or 1 mg/ml Neomycin. To create the reconstituted cell lines, HeLa GABARAPL2 KO cells (pLenti-Neo) were used and lentiviral transductions of HA-GABARAPL2 WT or mutants were performed, with a selection of 2 µg/ml of Puromycin or 1 mg/ml Neomycin.

CCCP was resuspended in DMSO and cells were treated with 40 μM for specific timepoints (30 min - 4h). Human IFNγ (AF-300-02; Peprotech) was added to cells for 24 hours at a final concentration of 10 ng/ml. Nutrient starvation was achieved by replacing DMEM with EBSS (Gibco) for 30 min - 4h. BafilomycinA1 was resuspended in DMSO and cells were treated with 200 nM for specific timepoints. MRT67307 (a specific TBK1 inhibitor) was used at 5 μM for 4 hours.

Plasmid transfections were performed with 3 μl GeneJuice (Merck Millipore), 0.5 μg plasmid DNA in 200 μl Opti-MEM (Life Technologies). After incubation for 15 min, the solution was added to the cells, which were lysed in lysis buffer or fixed with 4% paraformaldehyde 48 hours later.

SiRNA transfections were performed with 3 μl RNAiMax (Invitrogen), 20 nM siRNA (IRGQ #1: TTTGTGCTACCGGCGAACTG, IRGQ #2: GAATGCACTCAG-TAAGGGAA, IRGQ #3: CGTGAGGCCTTTGAGACCGG; TBK1 #1: 5′-GACAGAAGUUGUGAUCACATT-3′) all purchased from Sigma) in 150 μl Opti-MEM (Life Technologies). After incubation for 30 minutes, the solution was added to the cells cultured in a 6-well dish, which were lysed in lysis buffer or fixed with 4% paraformaldehyde 72 hours post transfection.

### Salmonella infections

Salmonella SL1344 (WT) were streaked out on LB plates and single colonies were picked to inoculate 2 ml of LB media (containing appropriate antibiotics and 0.3 M NaCl) to grow at 37°C for 16 hours. The overnight culture was then diluted 1:33 in LB media (containing appropriate antibiotics and 0.3M NaCl). After 2.5 hours, the OD600 of Salmonella was determined and the cells were infected for 30 min with a MOI of 150 (considering that OD600=1 has ∼1,3x10^9^ bacteria/ml). The infection media was then exchanged to DMEM (+ 10% FBS) with 50 μg/ml Gentamycin and cells were lysed or fixed after indicated time points.

### Immunofluorescence for confocal microscopy imaging

HeLa or U2OS cells were seeded onto glass coverslips in 12-well culture dishes and treated accordingly. Cells were washed in phosphate-buffered saline (PBS) before fixation with 4% paraformaldehyde for 15 minutes at room temperature. The coverslips were washed a further three times before permeabilization of the cells with 0.5% Triton X-100 in PBS for 10 minutes at room temperature. Cells were rinsed with PBS before being incubated for 1 hour in 1% bovine serum albumin (BSA) in PBS for 1 hour. Primary antibody incubation was done for 1 hour in a humidified chamber with 1% BSA in PBS.

After thorough washes in PBS, cells were incubated with secondary antibodies, 1% BSA in PBS for 1 hour in the dark. Cells were washed three more times in PBS and once with deionized water before being mounted onto glass slides using ProLong Gold mounting reagent (Life Technologies), which contained the nuclear stain 4′,6-diamidino-2-phenylindole (DAPI). Slides were imaged using a Leica microscope Confocal SP 80 fitted with a 60x oil-immersion lens.

### Immunofluorescence for Yokogawa CQ1 microscopy imaging

HeLa or T-REx mCherry-GFP GABARAP-L2 WT, S10A and S10D mutant U2OS cells were seeded onto black, clear flat bottom 24- or 96-well plates (2000 cells/well for 96-well plates and 18000 cells/well for 24-well plates). When indicated, cells were treated. Cells were washed in PBS before fixation with 4% paraformaldehyde for 15 minutes at room temperature. Cells were rinsed with PBS before being incubated for 1 hour in permeabilization and primary antibody solution (0.1% Saponin (47036; Sigma), 5 mM MgCl_2_, 5% BSA in PBS). After washes in PBS, cells were incubated with Hoechst 33342 (R37605; Thermo Fisher), Alexa Fluor 647 Phalloidin (#8940; Cell Signaling Technology) and Alexa Fluor secondary antibodies in antibody solution (0.1% Saponin (47036; Sigma), 5 mM MgCl_2_, 5% BSA in PBS) for 1 hour in the dark. Plates were imaged using a Yokogawa CQ1 microscope.

### Cell lysis

For lysis, cells were washed with PBS and scraped on ice in IP lysis buffer (50 mM Hepes, pH 7.5, 150 mM NaCl, 1 mM EDTA, 1 mM EGTA, 1% Triton X-100, 25 mM NaF, 5% glycerol, 10μM ZnCl2) or total cell lysis buffer (50 mM Tris HCl, pH 7.5, 1 mM EDTA, 1% SDS, 25 mM NaF, 1μl/ml Benzonase (71205-25KUN; Millipore)), both supplemented with complete protease inhibitors (cOmplete, EDTA-free; Roche Diagnostics) and phosphatase inhibitors (P5726, P0044; Sigma). Extracts were cleared by centrifugation at 15000 rpm for 15 minutes at 4°C.

### Immunoprecipitation of overexpressed proteins

Cleared cell extracts were mixed with HA-agarose beads (A2095; Sigma), Flag-M2 agarose beads (A2220; Sigma), RFP-Trap_A beads (rta-10; ChromoTek) or GFP-Trap_A beads (gta-10; ChromoTek) 16 hours at 4°C on a rotating platform. The beads were washed four times in IP lysis buffer. Immunoprecipitated and input samples were reduced in SDS sample buffer (50 mM Tris HCl, pH 6.8, 10% glycerol, 2% SDS, 0.02% bromophenol blue, 5% β-mercaptoethanol) and heated at 95°C for 5 minutes^33,34^.

### Immunoprecipitation of endogenous HA-GABARAPL2

HeLa cells stably expressing HA-tagged GABARAP-L2 were seeded in 15-cm dishes and cultured to ∼80% confluency prior to lysis. Cells were treated as indicated (e.g., siRNA knockdown, CCCP, or *Salmonella* infection). Cells were washed once with 10 mL PBS and lysed in 150 μL Total Cell Lysis (TCL) buffer (1% SDS, 50 mM Tris-HCl pH 7.5, 1 mM EDTA, 25 mM NaF, supplemented with protease inhibitor cocktails 2 and 3). Lysates were immediately transferred into 2 mL Eppendorf tubes, diluted with 1.35 mL of Normal Lysis (NL) buffer (150 mM NaCl, 50 mM HEPES pH 7.5, 1 mM EDTA, 1 mM EGTA, 10% (v/v) glycerol, 1% (v/v) Triton X-100, 10 μM ZnCl₂, 25 mM NaF, supplemented with protease inhibitor cocktails 2 and 3), and placed on ice. To degrade nucleic acids and reduce viscosity, 1 μL of Benzonase (Sigma-Aldrich) was added to each lysate, followed by incubation on ice for 20 minutes. Lysates were clarified by centrifugation at maximum speed for 10 minutes at 4 °C.

Input samples were prepared by transferring 50 μL of the cleared supernatant into a fresh tube, adding 10 μL of 4× SDS sample buffer, heating for 10 minutes at 95 °C, and storing at -20 °C.

HA immunoprecipitations were set up by incubating the remaining lysates with 20 μL of pre-equilibrated anti-HA magnetic beads (e.g., Thermo Fisher) overnight at 4 °C with gentle rotation in 1.5 mL tubes. The following day, beads were washed four times with chilled NL buffer. After the final wash, beads were dried using a 27G ¾’’ needle to minimize residual buffer volume. Proteins were eluted by adding 20 μL of 2× SDS sample buffer directly to the beads, followed by heating at 95 °C for 10 minutes.

### Protein binding assays

GST or GST-IRGQ were immobilized on glutathione-Sepharose beads (GE Healthcare) and combined with purified His-GABARAP-L2 in protein binding buffer (150 mM NaCl, 50 mM, Tris, pH 7.5, 0.1% Nonidet P-40, supplemented with 5 mM DTT and 0.25 mg/mL BSA). The proteins were incubated on a rotating platform at 4°C for 16 hours. After five washes with buffer, proteins were diluted with SDS sample buffer (62.5 mM Tris-HCl pH 6.8, 10% (v/v) glycerol, 2% (w/v) SDS, 0.02% (w/v) bromophenol blue, 5% (v/v) β-mercaptoethanol), resolved by SDS-PAGE and analyzed by immunoblotting with the indicated antibodies.

### Kinase assays

GABARAP-L2 WT and mutant proteins were incubated in 20 μl phosphorylation buffer (50 mM Tris HCl, pH 7.5, 10 mM MgCl2, 0.1 mM EGTA, 20 mM ß-glycerophosphate, 1 mM DTT, 0.1 mM Na_3_VO_4_, γ^P32^ ATP (500 cpm/pmol; SRP-201; Hartmann Analytic)) with 50 ng of recombinant GST-TBK1 for 15 minutes at 30°C. The kinase assay was stopped by adding SDS sample buffer containing 1% β-mercaptoethanol and heating at 95°C for 5 minutes. The samples were resolved by SDS-PAGE, and the gels were stained with InstantBlue (expedeon) and dried. The radioactivity was analysed by autoradiography^35^.

### Western blotting

For immunoblotting, proteins were resolved by SDS-PAGE and transferred to PVDF membranes. Blocking and primary antibody incubations were carried out in 5% BSA in TBS-T (150 mM NaCl, 20 mM Tris, pH 8.0, 0.1% Tween-20), secondary antibody incubations were carried out in 5% low-fat milk in TBS-T and washings in TBS-T. Blots were developed using Western Blotting Luminol Reagent (sc-2048; Santa Cruz). Immunoblot bands were quantified using ImageJ software. All Western blots shown are representative.

### Antibodies

The following antibodies were used in this study: anti-HA-tag (11867423001; Roche), anti-FlagM2-tag (F3165; Sigma), anti-GFP-tag (Living Colors 632592; Clontech), anti-His-tag (11922416001; Roche), anti-vinculin (V4505; Sigma), anti-TBK1 (#3013; Cell Signaling Technology), anti-pTBK1 (pS172; #5483; Cell Signaling Technology), anti-IRGQ (HPA043254; Sigma), anti-GAPDH (#2118; Cell Signaling Technology), anti-GABARAP-L2 (PM038; MBL), anti-LAMP1 (H4A3; DSHB), anti-p62 (M162-3; MBL), anti-IkBa (#9247; Cell Signaling Technology), anti-Histone H3 (ab1791, Abcam), anti-pSTAT1 (pY701; #7649; Cell Signaling Technology), anti-ATG3 (#3415; Cell Signaling Technology), anti-ATG7 (8558S, CST),anti-LC3B (PMO36; MBL), anti-ULK1 (8054S, CST), anti-MFN1 (14739S, CST), anti-pS10 GABARAP-L2 (was generated by immunGlobe^®^, a chemically synthesized peptide (GABARAP-L2 aa4-15) bearing a phosphate group at S10 (Ac-MFKEDH(pS)LEHRC-NH_2_) was used for immunization). Primary antibodies used for Western blotting were diluted 1:1000 and for immunofluorescence studies 1:200. Secondary HRP conjugated antibodies goat anti-mouse (sc-2031; Santa Cruz), goat anti-rabbit (sc-2030; Santa Cruz) and goat anti-rat (sc-2006; Santa Cruz) were used for immunoblotting. Anti-rat Alexa Fluor 647 (A-21247; Life Technologies), anti-rat Cy3 (712166153; Jackson Lab), anti-mouse Alexa Fluor 405 (A-31553; Life Technologies), anti-mouse Cy3 (715-165-151; Dianova), anti-mouse Alexa 647 (A-31626; Life Technologies) were used for immunofluorescence studies.

### Mass Spectrometry

Cells were treated, lysed and IP was performed as stated above, after which trypsin digestion and peptide desalting were performed. Digested peptides were acidified with trifluoroacetic acid (TFA) (Sigma Aldrich) to inhibit trypsin and to acidify peptides for SDB-RPS StageTip desalting. Acidified peptides were loaded onto the SDB-RPS StageTips and then washed with 0.1% (v/v) TFA. Peptides were eluted using a two-step elution with 0.1% (v/v) TFA, 80% (v/v) ACN and then dried using a speed-vacuum concentrator (30-45 min at 45-60°C). Dried peptides were stored at -20°C.

Peptides were analyzed on an Orbitrap EliteTM or Q Exactive HF mass spectrometer (ThermoFisher). The raw data was analyzed using MaxQuant 1.6.5.0 with standard settings and activated LFQ quantification. The database used to identify the peptides was the human reference protein database (uniprot downloaded December 2017) and the FDR was set to 1% on protein, PSM and site decoy level. Statistical analysis was done with Perseus. Proteins were defined as interactors, if they passed a 5% FDR corrected one sided two-sample T-test with a minimal enrichment factor of two.

### Phos-tagTM SDS-PAGE

Phos-tagTM acrylamide (Wako) gels were used as indicated by the supplier. Gels were prepared with 10% acrylamide, 50 μM phos-tagTM and 100 μM MnCl_2_. Cells were lysed in SDS sample buffer supplemented with 10 μM MnCl2.

### Proximity ligation assay (PLA)

HeLa cells were seeded onto black, clear flat bottom 96-well plates (2000 cells/well), treated with 10ng/ml IFNγ for 24 hours and infected with Salmonella. Cells were washed in PBS before fixation with 4% paraformaldehyde for 15 minutes at room temperature. Rabbit anti-IRGQ (HPA043254; Sigma) and mouse anti-HA (MMS-101P; Covance) were used with the respective Duolink in situ PLA probes (DUO92001, DUO92005; Sigma) and the PLA Duolink in situ detection reagent kit (DUO92008, Sigma) according to manufacturer’s instructions. After washes in PBS, cells were incubated with Hoechst 33342 and Alexa Fluor 647 Phalloidin for 1 hour in the dark. Plates were imaged using a Yokogawa CQ1 microscope and quantification of the PLA signal was performed using the Yokogawa CQ1 software.

### Protein expression and purification

GST or His-tagged fusion proteins were expressed in E. coli strain BL21 (DE3). Bacteria were cultured in LB medium supplemented with 100 μg/mL ampicillin at 37°C in a shaking incubator (150 rpm) until OD600 ∼0.5-0.6. Protein expression was induced by the addition of 0.5 mM IPTG and cells were incubated at 16°C for 16 hours. Bacteria were harvested by centrifugation (4000 rpm) and lysed by sonication in GST lysis buffer (20 mM Tris HCl, pH 7.5, 10 mM EDTA, pH 8.0, 5 mM EGTA, 150 mM NaCl, 0.1% β-mercaptoethanol, 1 mM PMSF) or His lysis buffer (25 mM Tris HCl, pH 7.5, 200 mM NaCl, 0.1% β-mercaptoethanol, 1 mM PMSF, 1mg/ml lysozyme). Lysates were cleared by centrifugation (10000 rpm), 0.05% of Triton X-100 was added and the lysates were incubated with glutathione Sepharose 4B beads (GE Life Sciences) or Ni-NTA agarose beads (Thermo Fisher) on a rotating platform at 4°C for 1 hour. The beads were washed five times either in GST wash buffer (20 mM Tris HCl, pH 7.5, 10 mM EDTA, pH 8.0, 150 mM NaCl, 0.5% Triton X-100, 0.1% β-mercaptoethanol, 1 mM PMSF) or His wash buffer (25 mM Tris HCl, pH 7.5, 200 mM NaCl, 0.05% Triton X-100, 10 mM Imidazole). The immobilized proteins were reconstituted in GST storage buffer (20 mM Tris HCl, pH 7.5, 0.1% NaN3, 0.1% β-mercaptoethanol) or eluted with His elution buffer (25 mM Tris HCl, pH 7.5, 200 mM NaCl, 300 mM Imidazole) and dialysed in (25 mM Tris HCl, pH 7.5, 200 mM NaCl) at 4°C for 16 hours. Recombinant GST-TBK1 was obtained from the MRC PPU DSTT in Dundee, UK (#DU12469)^18^.

### FACS

T-REx mCherry-GFP GABARAP-L2 WT, S10A and S10D mutant U2OS were cultivated in cell culture flasks until ∼80 % confluency and transferred to 6-well plates (500.000 cells/well) for the experiments (3 replicates/condition for every cell line). Cells were treated over night with doxycycline (final concentration: 1 µg/ml) to induce mCherry-GFP expression. Negative control was not induced. After induction, cells were washed once with PBS and treated as follows: untreated (stopped after 2 h), CCCP (40 µM, 2 h), EBSS (6 h) and EBSS + Bafilomycin (200 nM, 6 h). After treatments, cells were detached using trypsin/EDTA, centrifuged (5 min, 500 x g) and resuspended in 200 µl of FACS buffer (PBS + EDTA + FBS). FACS was performed directly at BD Symphony A5. Following singlet gating, cells were gated for high mCherry^+^ GFP^+^ cells using FlowJo software (version 10). The mean fluorescence intensity (MFI) for mCherry and GFP was calculated for each sample. Fold changes were calculated and normalized to the mean of each untreated cell line (WT, S10A and S10D, respectively). Analysis was performed using GraphPad Prism software.

### Modelling and simulations of IRGQ-ATG8 complexes

Full-length sequences of IRGQ (UniProtKB: Q8WZA9), GBRL2 (UniProtKB: P60520), MLP3B (LC3B, UniProtKB: Q9GZQ8), and ATG7 (UniProtKB: O95352) were used to model the 3D structures of various complexes using AlphaFold2-multimer model (AF2) multimers: (1) IRGQ-GBRL2, (2) IRGQ-GBRL2-ATG7, and (3) IRGQ-GBRL2-LC3B-ATG7. We obtained 100 models for each complex using the default AFv2.0 parameters (database updated, 2025). To evaluate the stability of the IRGQ-GBRL2 complex, we performed all-atom molecular dynamics (MD) simulations using the PDB structure of the IRGQ_1-189_-GABARAPL2 complex (PDB ID: 8Q6Q) as the initial model. Using the CHARMM-GUI server^36^, we refined the native (WT) IRGQ-GBRL2 complex and also modeled the phosphorylated version (S10-PO4). We ensured that the disulfide bridge between C152 and C158 was preserved in our models. Protein complexes were placed in an octahedral box and solvated with the TIP3P water model and physiological salt concentration (150 mM NaCl). All-atom MD simulations of the IRGQ-GBRL2 complexes were performed with GROMACS (v 2021.5)^37^ using the CHARMM36m force field. Initially, the system was minimized using the steepest-descent algorithm until the maximum force reached 1000 kJ nm^-1^. The equilibration phase was run in an NVT ensemble with the v-rescale thermostat at 310 K (*τ*_T_ = 1 ps)^38^. Position restraints were applied to the backbone (400 kJ mol^-1^ nm^-2^) and sidechain (40 kJ mol^-1^ nm^-2^) atoms to equilibrate the water. During the production run, the pressure was maintained at 1 bar (τ_P_ = 5 ps, compressibility = 4.5E-05) using an isotropic c-rescale barostat^39^. Three replicates of production runs were simulated in each condition for 1000 ns with a 2-fs timestep. Coarse-grained MD simulations for the IRGQ-GBRL2 complexes were performed using (WT) and a phosphomimetic variant (S10D). The Martini force field was used to map the atomistic structure of the top-ranked AF2 model with the martinize2.py script^40^. For both variants, we employed the Go-Martini 3.0 model. Secondary structure assignments were done using DSSP, followed by automatic identification of disulfide bonds. Backbone restraints were applied with a force constant of 1000 kJ mol^-1^ nm^-2^, Go-like native contacts were modeled (*ε* = 12, residue distance cutoff = 3), along with predefined intrinsically disordered regions (regions 1-7, 179-191, 331-430, and 617-623) explicitly treated to preserve their conformational flexibility. The CG models were placed in a hexagonal simulation box, solvated with coarse-grained water beads, with 0.15 M NaCl. MD simulations were performed using GROMACS (v 2021.5)^37^. Systems were first energy minimized for 3000 steps with the steepest-descent approach, followed by an equilibration in an NPT ensemble at 310 K. Temperature and pressure were maintained, respectively, with a v-rescale thermostat (*τ*_T_ = 1 ps) and an isotropic c-rescale barostat (*τ*_P_ = 5 ps, compressibility = 4.5E-05). During the production runs, we used the Parrinello-Rahman scheme (*τ*_P_ = 12 ps)^41^ to maintain system pressure. Simulations were performed with a 20-fs time step for 1000 ns, and 15 replicates were run for each system. Distances and contact maps for pairwise residue-residue interactions within each complex were computed using in-house scripts based on MDAnalysis v2.9.0^42^. Contact maps were computed by counting pairwise residue contacts between chain A and chain B according to AB_cnts_ = [Σ_i∈A_Σ_j∈B_ σ|r_ij_|)], where the sums extend over heavy atom positions of interacting residues (ij) and σ(|r_ij_| = 1 - [0.5 - 0.5(tanh((|r_ij_|-a/b))]), a smooth sigmoidal counting function to limit interactions below the cut-off distance (r_ij_ ≤ a), where the cutoff parameter, a was set to 5 Å for atomistic simulations and 10 Å for CG simulations, while the smoothing parameter *b* was set to 0.5 and 1.0, respectively. Binding free energies for the WT and S10PO4 variant complexe of atomistic simulations were estimated using the MMPBSA approach as implemented in the gmx_MMPBSA program with default parameters^43^. The binding free energy was decomposed into gas-phase and solvation contributions, with the gas-phase term (Δ*G*_gas_) comprising van der Waals and electrostatic energies, and the solvation term (Δ*G*_solv_) including polar electrostatic contributions computed using the Poisson-Boltzmann model and nonpolar contributions estimated from solvent-accessible surface area. Per-residue free-energy decomposition was performed for both WT and S10PO4 systems to quantify residue-specific energetic contributions, and uncertainties were estimated from variability across independent replicas.

### Statistical analysis

All experiments have a minimum of three biological replicates. Yokogawa experiments additionally have six technical repeats for each biological repeat. Data are presented as the mean with error bars indicating the s.d. (standard deviation). Statistical significance of differences between experimental groups was assessed with Student’s T-test. Differences in means were considered significant if p < 0.05. Differences with p < 0.05 are annotated as *, p < 0.01 are annotated as ** and p < 0.001 are annotated as ***. All western blots shown are representative of biological replicates. Immunofluorescent images were analyzed with CellProfiler 4.2.8.

### Sequence alignments

Sequence alignments were performed using the Clustal algorithm^44^ with Ensemble identifiers. This approach allowed for the rapid and accurate alignment of protein sequences from the Ensemble database, facilitating the identification of conserved motifs and regions of interest.

## Data availability

All data are available in the main manuscript text or the supplementary materials and will be made available upon request. The mass spectrometry proteomic data have been deposited to the ProteomeXchange Consortium via the PRIDE partner repository with the following dataset identifiers: IRGQ-GABARAPL2 complex interactome data - PXD066671, GABARAPL2 interactome - PXD066665.

## Author contributions

U.G.-M. designed/performed most of the experiments and analyzed the data. L.H., and I.D., conceived the study. S. A. P-C., and A.C., performed molecular modeling and simulations with support and supervision from R.M.B. L.H., P.L, B. C., M. A. and A.V.A., performed some experiments and analyzed data. L.H. and I.D. managed the project and supervised experiments. U.G.-M. and L.H. wrote the manuscript with input from all authors.

## Acknowledgments

We thank Prof Kraiczy for the provided S2 Salmonella working space. We are also grateful to Henry Bailey for valuable discussions and providing critical data to our previous manuscript^16^. We acknowledge all members of the Quantitative Proteomics Unit at IBC2 (Goethe University, Frankfurt), in particular Thorsten Mosler and Georg Tascher, for support and expertise in proteomics methodology and data analysis, Martin Adrian-Allgood and Julia Pomirska for technical help and measurements, Kristina Wagner for preparing LC columns, and David Krause for help in (bio)informatics. We thank the FACS facility at the Helmholtz-Zentrum für Infektionsforschung GmbH, Braunschweig. We thank the Deutsche Forschungsgemeinschaft (German Research Foundation, DFG) for funding the liquid chromatography (LC)-MS system (easy nLC1200, Orbitrap Fusion LUMOS) used in this study (FuGG Project-ID: 403765277). The research was funded by Dr. RolfM. Schweite Stiftung to I.D. (project 13/2017), grants from the Goethe University Frankfurt to L.H. (Nachwuchswissenschaftler grant 710000624), and GRADE A/B Focus to L.H. (PID003790), as well as the LOEWE Zentrum Frankfurt Cancer Institute Discovery & Development Grant to L.H. (21001366). S.A.P-C, A.C., and R.M.B. thank the Center for Supercomputing, Goethe University Frankfurt (GUF), for computing time on the Goethe-HLR cluster. Additionally, this work was supported by the Clusterproject ENABLE funded by the Hessian Ministry for Science and the Arts to R.M.B, and L.H.; CRC project on selective autophagy, grant/award number: project-ID 259130777 to R.M.B, I.D. and L.H.; the Leistungszentrum Innovative Therapeutics (TheraNova) funded by the Fraunhofer Society and the Hessian Ministry of Science and Arts to I.D. and U.G.M. Work of the Herhaus lab at the Helmholtz-Zentrum für Infektionsforschung GmbH, Braunschweig is supported by the by the Microbial Stargazing program of the German Federal Ministry of Education and Research (BMFRT) under grant number 01KX2324. Responsibility for the content of this publication lies with the author.

---

**Figure S1:**
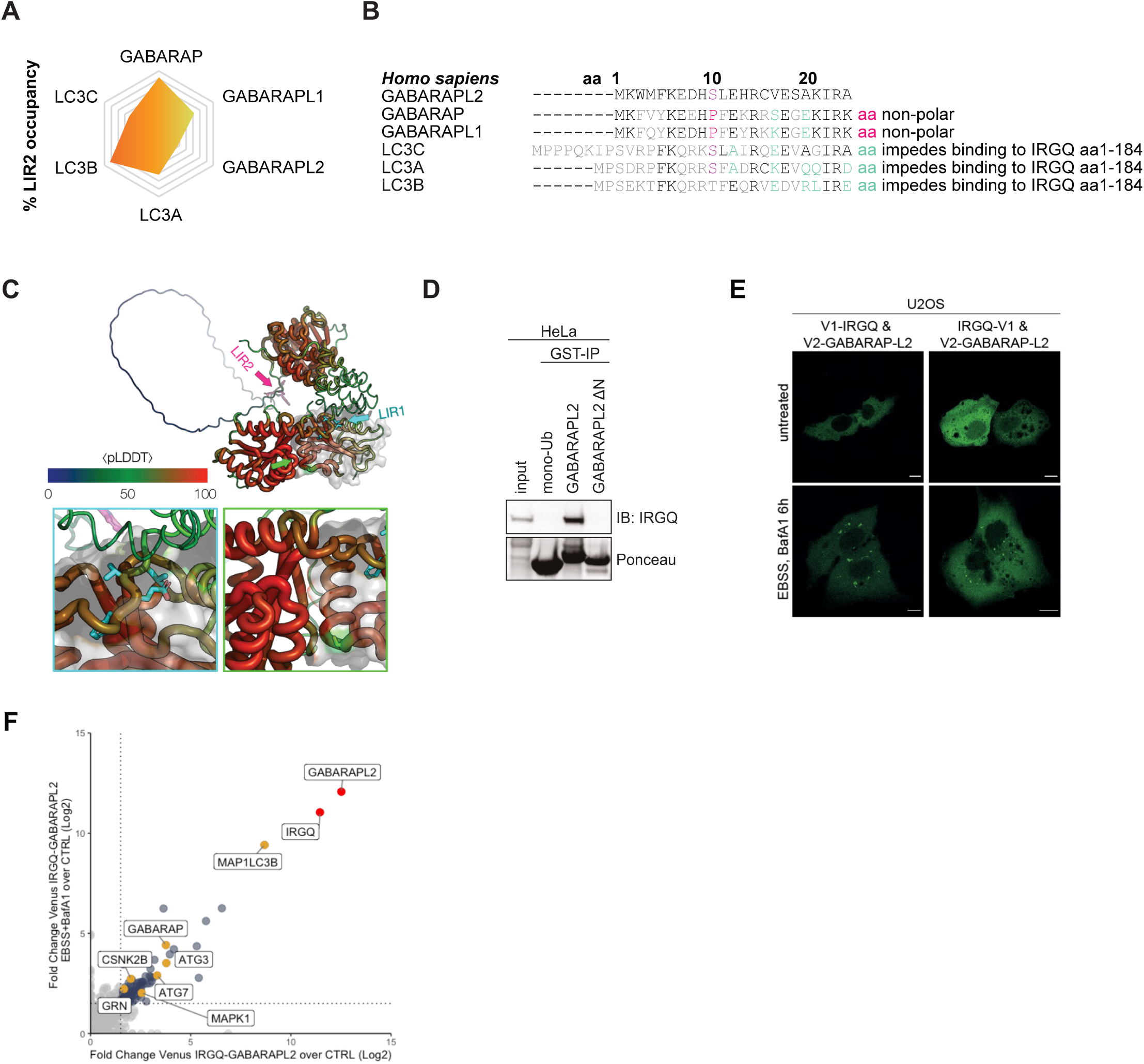
**(A)** Percentage of LIR2-LDS binding mode as the predicted complex between IRGQ-hATG8 in the 25 top-ranked AlphaFold2-multimer models from previous study^16^. **(B)** Sequence alignment of hATG8 N-terminals highlighting amino acids that would impede binding to IRGQ, in red polar residues, in green residues potentially causing steric clashes. **(C)** Top-ranked AlphaFold2 model for the IRGQ-GABARAPL2 complex. Confidence on local predicted structure (pLDDT) is averaged over top 100 models and mapped onto the structure (rainbow) GABARAPL2 is represented as a gray surface. Arrows indicate three key regions of IRGQ: LIR1 (cyan), LIR2 (pink), and switch II (green). Bottom panels show zoom-up of the LIR-LDS (left) and switch I/II protein-protein interfaces (right, ⟨pLDDT⟩ > 80). **(D)** SDS-PAGE and Western blot of an *in vitro* GST-pulldown using purified GST-mono-ubiquitin, GST-GABARAPL2 or GST-GABARAPL2 ΔN incubated with HeLa cell extract. **(E)** Immunofluorescence of U2OS cells, transiently expressing Vn-IRGQ and Vn-GABARAPL2, or IRGQ-Vn and GABARAPL2-Vc. Autophagy was induced by the addition of EBSS for 6 hours and lysosomal degradation was blocked by BafA1 (200 nM) treatment. **(F)** Scatter plot representing the Student’s T-test difference from Vn-IRGQ-Vc-GABARAPL2 over VnVc-ORF IPs in Basal conditions and the Student’s T-test difference from Vn-IRGQ-Vc-GABARAPL2 over VnVc-ORF Ips upon autophagy induction. The baits IRGQ and GABARAPL2 are marked in red, significant interaction partners in blue and autophagy-related proteins marked in yellow.

**Figure S2:**
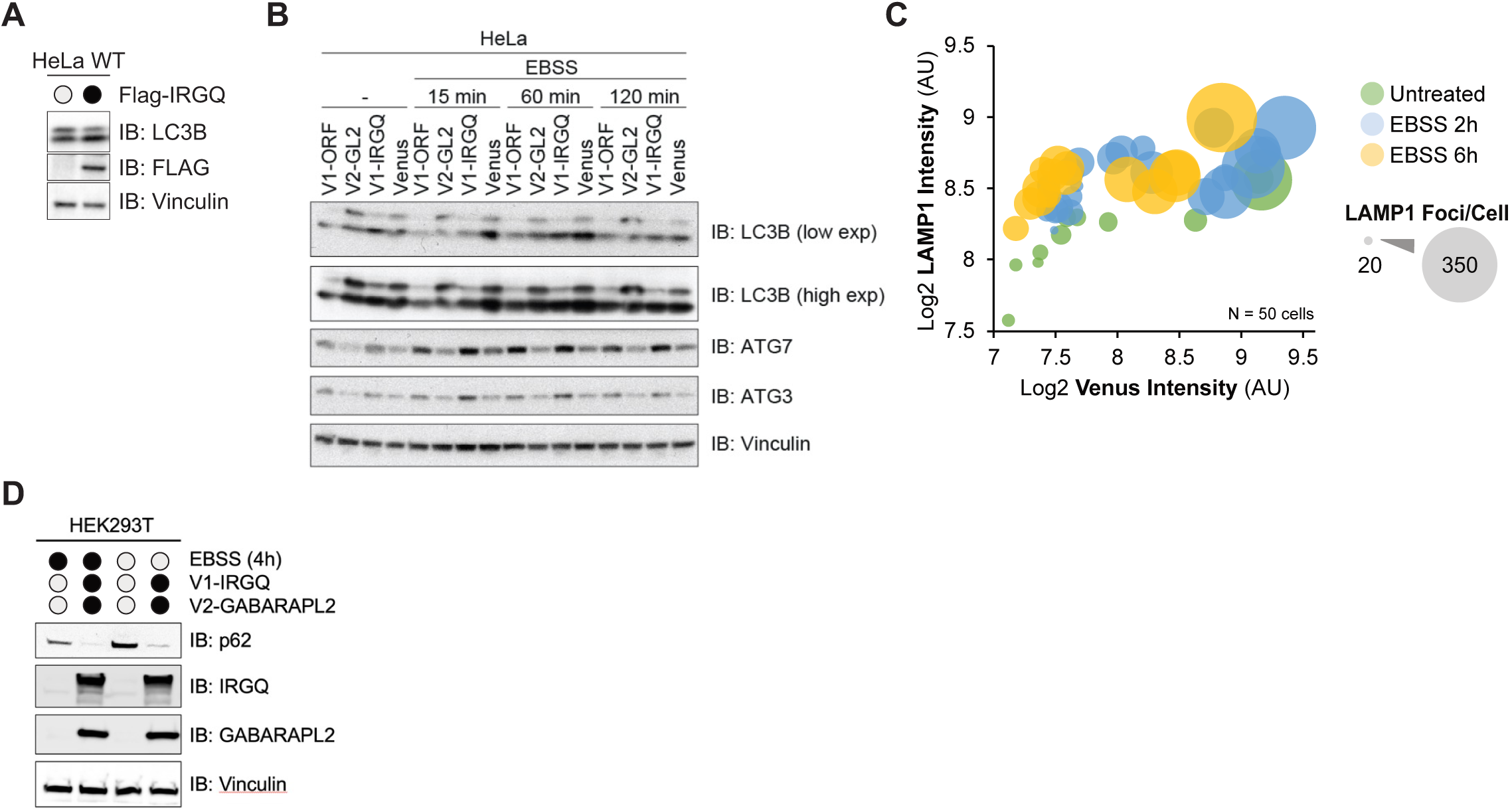
**(A)** SDS-PAGE and Western blot of HeLa cell lysates expressing transfected Flag-IRGQ. **(B)** SDS-PAGE and Western blot of HeLa cell lysates stably expressing V1-ORF, V2-GABARAPL2, V1-IRGQ or V1-IRGQ & V2-GABARAPL2 (Venus). Cells were either left untreated or treated with EBSS for indicated time points. **(C)** Scatter plot of ImageJ quantification from (D). Single cells were annotated as ROIs and intensities and puncta for LAMP1 and Venus were measured. RawIntDen values are plotted as Log2. Colors indicate the treatment and size of bubbles indicate the number of LAMP1 puncta per cell; n=50 cells. **(D)** SDS-PAGE and Western blot of HEK293T cell lysates with transfected V1-IRGQ and V2-GABARAPL2 untreated or treated with EBSS (4h).

**Figure S3:**
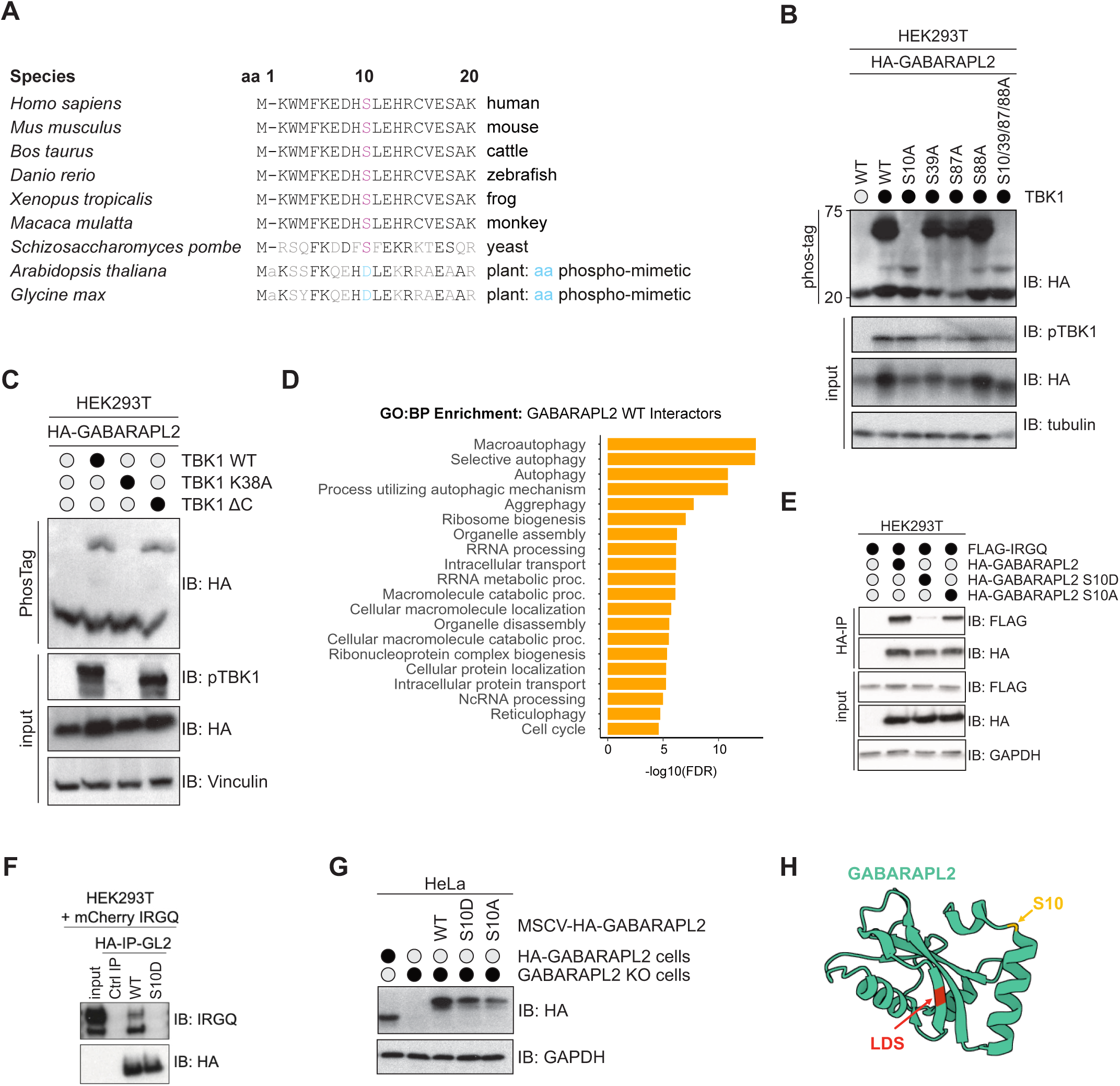
**(A)** Sequence alignment from multiple species for GABARAPL2 N-terminals (aa1-20) highlighted in red S10 conserved among the comparisons and in blue diverted amino acids in plants (D, phosphomimetic). **(B)** SDS-PAGE and Western blot of phos-tag gel with HEK293T cell lysates. Cells were transfected with TBK1 and HA-GABARAPL2 WT and mutants. **(C)** SDS-PAGE and Western blot of phos-tag gel with HEK293T cell lysates. Cells were transfected with HA-GABARA-L2 WT and TBK1 WT, TBK1 K38A (kinase inactive) and TBK1 ΔC-terminal coiled-coil region. **(D)** Gene Ontology enrichment of Biological Processes for HA-GABARAPL2 WT interactors, enrichment done with ShinyGO 0.77. **(E)** SDS-PAGE and Western blot of HEK293T cell lysates and HA-IPs. Cells were transfected with Flag-IRGQ, HA-GABARAPL2 WT, S10D or S10A and lysates used for HA-IPs. **(F)** SDS-PAGE and Western blot of HEK293T cell lysates and HA-IPs. Cells were transfected with mCherry-IRGQ and HA-GABARAPL2 WT or S10D and lysates used for HA-IPs. **(G)** SDS-PAGE and Western blot of HeLa GABARAP-L2 KO cells reconstituted with WT, S10A or S10D HA-GABARAPL2. **(H)** Crystal structure of GABARAPL2 (PDB:4CO7) with S10 marked in yellow and the LIR docking site (LDS) marked in red.

**Figure S4:**
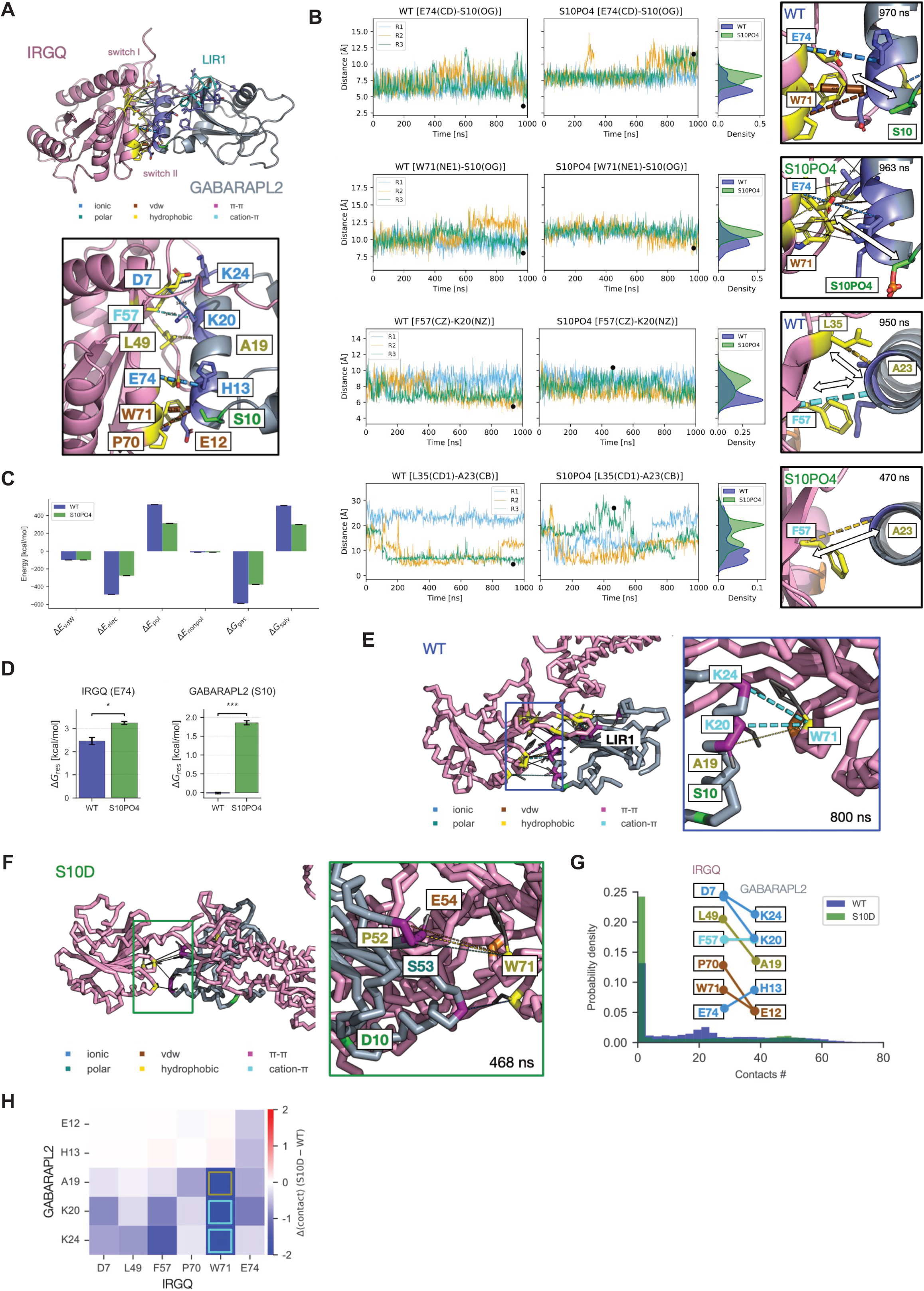
**(A)** Xray model (8Q6Q) of the N-terminal domain (pink) of IRGQ containing LIR1 (186-189, cyan) in complex with GABARAPL2 (grey). Helix 2 of GBRL2 (purple) makes contacts within a pocket formed by IRGQ switch I (partially disordered), switch II, and the linker spanning LIR1 site and the N-terminal G-domain (right, zoom-up). Key ionic interactions across this interface (>10 residue pair-wise contacts) between IRGQ1-186 (yellow) and GBRL2 (purple) contribute to the complex stability. **(B)** Phosphorylation of GABARAPL2 destabilizes the IRGQ-GBRL2 switch I and II interface. Time series of distances between interacting residue pairs estimated from atomistic MD simulations (3 replicates x 1000 ns) of WT complex (left), and the phosphorylated complex (middle, S10-PO4). Comparison of density distributions of distances (WT vs. S10-PO4) across select interface residue-pairs (top to bottom) for E74-S10(OG), W71(NE1)-S10(OG), F57(CZ)-K20(NZ), and L35(CD1)-A23(CB). (right) Selected snapshots (indicated times) corresponding of the complex simulations showing the structural rearrangements at the interface for S10-PO4 indicating destabilization. Distances are shorter in WT complex in comparison to S10-PO4. **(C)** MMPBSA energy computation and its decomposition into various terms shows relative contributions to the total binding energy of the IRGQ-GBRL2 complex (WT, blue; S10PO4, green). S10-PO4 complex displays an altered balance between electrostatic and solvation components, consistent with the structural reorganization of the interface induced by excess negative charge. **(D)** Residue-wise energy decomposition shows destabilizing effects. *** p < 0.001; * p < 0.05. **(E,F)** Representative snapshots from coarse-grained Go-Martini 3.0 simulations of IRGQ-GABARAPL2 complex (WT vs. S10D). Representative snapshot from WT complex simulations (15 replicates, 1000 ns each) showing preservation of the native interface. The LIR1 region remains stably engaged with GABARAPL2, and key interfacial residues (including K24, K20, A19, S10, and W71) maintain persistent contacts (right, zoom-up). By contrast, the S10D complex exhibits a disrupted native interface. The presumed interface residues mediate interactions with alternative sites on GABARAPL2 causing instability. **(G)** Distributions of #. of interface contacts in WT (blue) exhibits consistently higher interactions in comparison to S10D (green). **(H)** Heatmap showing the difference in pairwise residue-residue contact averages between WT and S10D (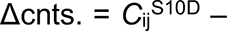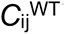). Most interface residue pairs strongly interact in WT complex in comparison to S10D consistent with altered electrostatic effects even in CGMD runs.

**Figure S5:**
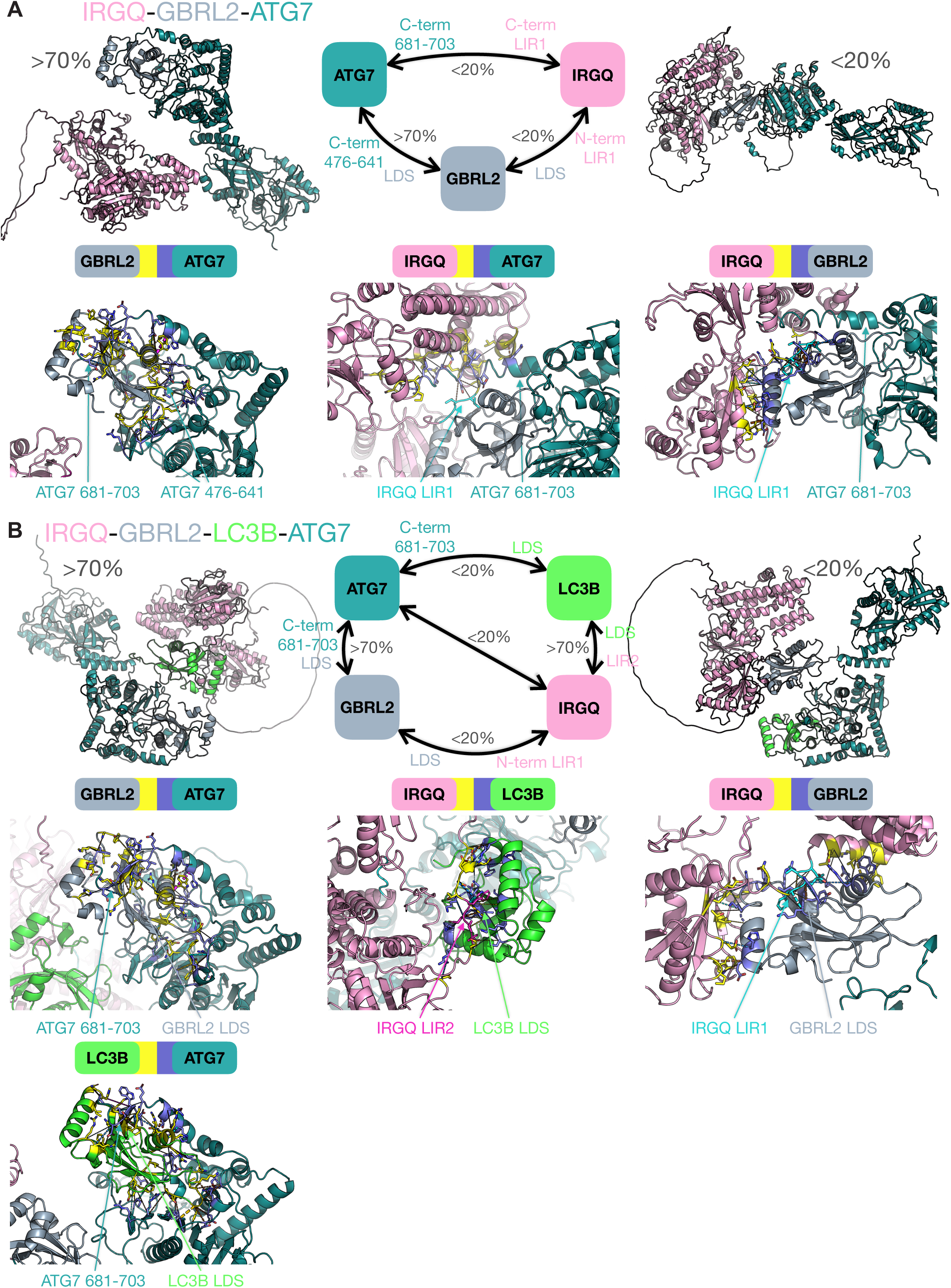
**(A)** AF2 multimers show differential IRGQ-hATG8 binding modes. Predicted IRGQ-GBRL2-ATG7 complexes display ATG7 C-terminal interacting with GBRL2 in > 70% of the models (top left), while < 20% of the models represent IRGQ and ATG7 interacting with GBRL2 using different interfaces (top right). Bottom panels display the pairwise interfaces (interfacing regions are coloured in yellow and violet for the first and second protein). **(B)** Predicted IRGQ-GBRL2-LC3B-ATG7 complexes display > 70% of models in which ATG7 interacts with GBRL2 and IRGQ interacts with LC3B (top left), while < 20% of the models present exchanged interactions, with ATG7 interacting with LC3B and IRGQ interacting with GBRL2 (top right). Bottom panels display the pairwise interfaces (interfacing regions are coloured in yellow and violet for the first and second protein).

**Figure S6:**
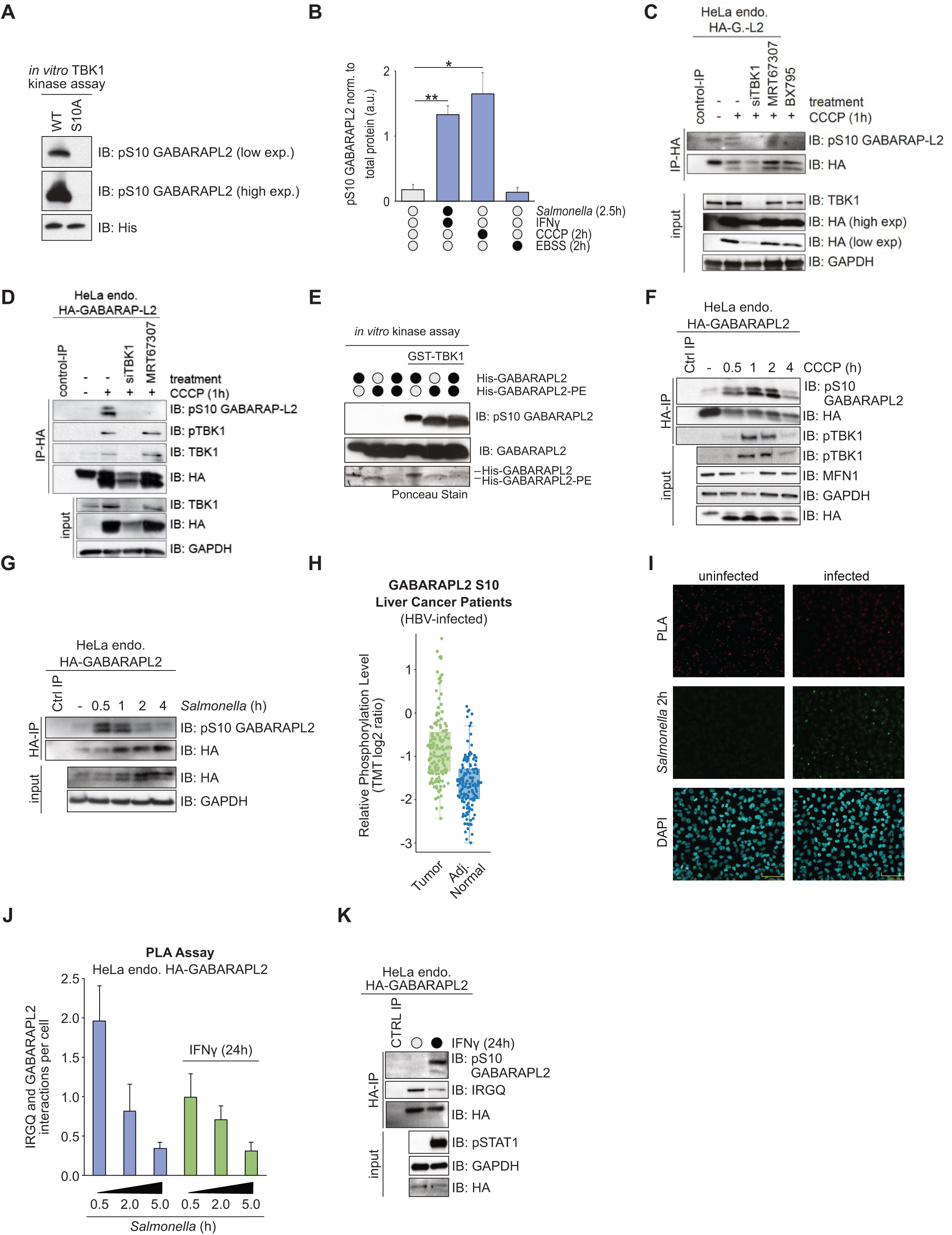
**(A)** SDS-PAGE and Western blot of *in vitro* TBK1 kinase assay with His-GABARAPL2 WT or S10A as substrate to test the pS10 GABARAPL2 antibody specificity. **(B)** ImageJ quantification from (4A) of pS10 GABARAPL2, normalized to total GABARAPL2 protein. Data are presented as the mean with error bars indicating the s.d. Statistical significance of differences between experimental groups was assessed with Student’s T-test. Differences with p<0.05 are annotated as * and p<0.01 are annotated as **; n=3. **(C)** SDS-PAGE and Western blot of HA-IPs from endogenously tagged HA-GABARAPL2 WT HeLa cells after treatment with TBK1 inhibitors (MRT67307 or BX795) or TBK1 siRNA knock-down, plus CCCP treatment (1h). **(D)** SDS-PAGE and Western blot of HA-IPs from endogenously tagged HA-GABARAPL2 WT HeLa cells after treatment with TBK1 inhibitor (MRT67307) or TBK1 siRNA knock-down, plus CCCP treatment (1h). **(E)** SDS-PAGE and Western blot of *in vitro* TBK1 kinase assay with His-GABARAPL2 and His-GABARAPL2-PE as substrate. **(F)** SDS-PAGE and Western blot of endogenously tagged HA-GABARAPL2 cells treated with 40 uM CCCP at specified timepoints. Lysates were used for HA-IP. **(G)** SDS-PAGE and Western blot of endogenously tagged HA-GABARAPL2 cells Infected with Salmonella (MOI:150) at specified timepoints. Lysates were used for HA-IP. **(H)** Relative Phosphorylation Level of GABARAPL2 S10 in HBV-derived liver samples from human samples. Data retrieved from www.cprosite.ccr.cancer.gov,^45^. **(I)** Images of PLA signal (red) from endogenous GABARAP-L2 and endogenous IRGQ. HeLa endogenous HA-GABARAPL2 cells were infected with *Salmonella* (GFP) for 2 hours and fixed cells were probed with the Duolink in situ PLA assay. HA and IRGQ only antibodies were used as negative controls to determine the background. **(J)** Yokogawa CQ1 quantification of average Duolink PLA signal from endogenous GABARAP-L2 and endogenous IRGQ. HeLa endogenous HA-GABARAPL2 cells were treated with 10 ng/ml IFNγ (24 hours), infected with Salmonella for indicated time points and fixed cells were probed with the Duolink in situ PLA assay. Data are presented as the mean with error bars indicating the s.d.; n=3; >3000 cells/condition. **(K)** SDS-PAGE and Western blot of endogenously tagged HA-GABARAPL2 cells treated with 10 ng/ml IFNγ (24 hours). Lysates were used for HA-IP.

**Figure S7:**
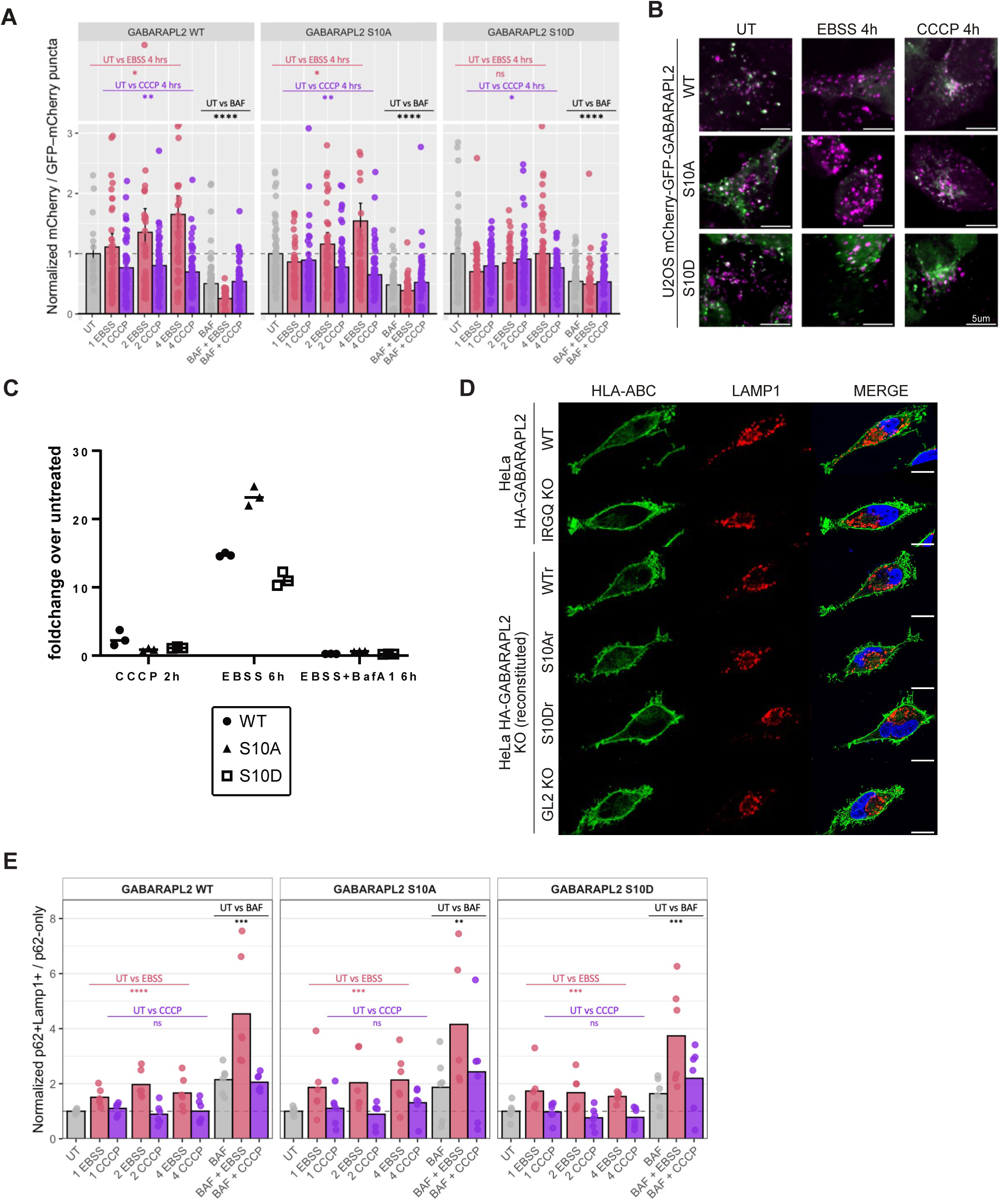
**(A)** GABARAPL2-to-lysosome flux under stress is lost for the S10D phosphorylation mutant. Quantification of GABARAPL2-to-lysosome flux in cells expressing GABARAPL2 WT, S10A, or S10D. Each point represents the mean mCherry/GFP-mCherry puncta ratio per well and per biological replicate (four independent experiments, two wells per condition). Data were first normalized to the mean untreated (UT) signal averaged across WT, S10A, and S10D within each replicate, and then rescaled so that UT = 1 for all genotypes in the final plot. Conditions correspond to nutrient starvation (EBSS; 1-4 h), mitochondrial depolarization (40 µM CCCP; 1-4 h), or lysosomal inhibition (200 nM BAF for 4 h; ± EBSS or CCCP). One-sample Wilcoxon signed-rank with Holm p-value correction. **** p < 0.0001; *** p < 0.001; ** p < 0.01; * p < 0.05; n.s., not significant. **(B)** Immunofluorescence of cells expressing mCherry/GFP-GABARAPL2 WT, S10A, or S10D. Conditions correspond to nutrient starvation (EBSS; 1-4 h), mitochondrial depolarization (40 µM CCCP; 1-4 h), or lysosomal inhibition (200 nM BAF for 4 h; ± EBSS or CCCP). **(C)** FACS analysis of of GABARAPL2-to-lysosome flux in cells expressing mCherry-GFP GABARAPL2 WT, S10A, or S10D. Each point represents the mean mCherry/GFP–mCherry puncta ratio per well and per biological replicate. Data is presented as fold change and normalized to the mean untreated cell line (WT, S10A or S10D, respectively). Conditions correspond to nutrient starvation (EBSS 6 h), mitochondrial depolarization (40 µM CCCP 2 h), or lysosomal inhibition (200 nM BAF for 6 h + EBSS 6 h). n=3. **(D)** Immunofluorescence of HeLa GABARAPL2 WT, IRGQ KO or GABARAPL2 KO cells reconstituted with WT, S10A or S10D HA-GABARAPL2. Autophagy was induced by the addition of EBSS for 4 hours and Bafilomycin A1 (200 nM). Fixed cells were probed with endogenous LAMP1 and HLA-ABC antibodies. Scale bar: 10 µm. **(E)** p62-to-lysosome flux under stress conditions is independent of GABARAPL2 S10 phosphorylation. Quantification of p62-to-lysosome flux in GABARAPL2 WT, S10A, and S10D cells. Each point represents the normalized mean ratio per well and per biological repeat (three independent experiments, two wells per condition). Values were normalized to the untreated (UT) mean within each genotype and repeat. Conditions represent starvation (EBSS, 1-4 h), mitochondrial depolarization (40 µM CCCP, 1-4 h), or lysosomal inhibition (200 nM BAF for 4 h; ± EBSS or CCCP). Grouped one-sample t-tests compared combined EBSS (E1-E4), CCCP (C1-C4), and BAF (BAF, EBSS + BAF, CCCP + BAF) groups to UT (= 1). **** p < 0.0001; *** p < 0.001; ** p < 0.01; * p < 0.05; n.s., not significant.

